# The interkingdom horizontal gene transfer in 44 early diverging fungi boosted their metabolic, adaptive and immune capabilities

**DOI:** 10.1101/2021.12.02.471044

**Authors:** Michał Aleksander Ciach, Julia Pawłowska, Paweł Górecki, Anna Muszewska

## Abstract

Numerous studies have been devoted to individual cases of horizontally acquired genes in fungi. It has been shown that such genes expand the hosts’ metabolic capabilities and contribute to their adaptations as parasites or symbionts. Some studies have provided an extensive characterization of the horizontal gene transfer (HGT) in Dikarya. However, in the early diverging fungi (EDF), a similar characterization is still missing. In order to fill this gap, we have designed a computational pipeline to obtain a statistical sample of reliable HGT events with a possibly minimal number of false detections. We have analyzed 44 EDF proteomes and identified 829 xenologs in fungi ranging from *Chytridiomycota* and *Blastocladiomycota* to *Mucoromycota*. We have identified several patterns and statistical properties of EDF HGT. Ancestrally aquatic fungi are generally more likely to acquire foreign genetic material than terrestrial ones. Endosymbiotic bacteria can be a source of useful xenologs, as exemplified by NOD-like receptors transferred to *Mortierellomycota*. Closely related fungi have similar rates of intronization of xenologs. The number of post-transfer paralogs of a protein can be described by a heavy-tailed Yule-Simons distribution. Post-transfer gene fusions complicate the landscape of HGT. We have designed a methodology to obtain a reliable, statistical sample of inter-kingdom xenologs across the tree of life of EDF to give a preliminary characterization of their general properties and patterns. We show that HGT is driven by bursts of gene exchange and duplication, resulting in highly divergent numbers and molecular properties of xenologs between fungal lineages. A close ecological relationship with another organism seems to be a predisposing condition for HGT, but does not always result in an extensive gene exchange. We argue that there is no universal approach for HGT identification and inter- and intra kingdom transfers require tailored identification methods. Our results help to better understand how and to what extent HGT has shaped the metabolic, adaptive, and immune capabilities of fungi.

## Introduction

Horizontal gene transfer (HGT) is a process of exchange of genetic material between distantly related species. It has been extensively studied in prokaryotes and proved to be an important mechanism that increases genomic diversity, contributes to niche adaptation, antibiotic resistance, new metabolic pathways, microbiome stability (Coyte et al. 2022) and numerous other traits. In eukaryotes, the significance and frequency of HGT remain mostly unclear, and the mechanisms of gene transfer beyond *Agrobacterium*-mediated gene transformation are still a matter of debate (Naranjo-Ortiz and Gabaldón 2020). It has been shown, however, that HGT does occur both within eukaryotes (including animals) and between eukaryotes and prokaryotes (Gabaldón 2020; Keeling and Palmer 2008).

Fungi represent very diverse life strategies, making them a unique model to study the connections between HGT and ecological adaptations. Genomes of fungi are protected against the incorporation of foreign DNA at multiple levels: starting from the physical separation of chromatin from the rest of the cell by the nuclear envelope, to foreign DNA elimination mechanisms RNAi and RIP, which are ubiquitous in fungi (Krassowski et al. 2018; Clutterbuck and John Clutterbuck 2011; Torres-Martínez and Ruiz-Vázquez 2017). On the other hand, the intake of foreign DNA may be facilitated by hyphal fusion (Priest, Yadav, and Heitman 2020; Feurtey and Stukenbrock 2018; Richards and Talbot 2013). Although classical theorists link eukaryotic HGT with phagocytosis (Doolittle 1998). The complex interplay of factors facilitating and preventing the incorporation of foreign DNA places fungi somewhere between bacteria and animals in terms of the rate of HGT and contributes to large differences in the numbers of HGT between different fungal lineages.

Several horizontally transferred genes have documented adaptive roles for fungal fitness. A fungal pathogen of insects, *Metarhizium robertsii*, has acquired sterol carrier and peptidase genes that facilitate the penetration of the victim’s body (Zhao et al. 2014; Zhang et al. 2019). Multiple transfers of toxins in plant-infecting fungi have improved their ability to infect new hosts (Mehrabi et al. 2011; Ambrose, Koppenhöfer, and Belanger 2014; B.-F. Sun et al. 2013). Bacteria have been described as a source of extensive HGT in plant-associated *Colletotrichum* sp. (Jaramillo, Sukno, and Thon 2015), *Verticillium dahliae* (Shi-Kunne et al. 2019) and animal skin pathogens *Malassezia* (Ianiri et al. 2020). Spectacular examples of transfers of clusters of functionally-linked genes (Wisecaver and Rokas 2015; Wisecaver, Slot, and Rokas 2014), genes facilitating metabolism expansions (Gao et al. 2019) or even conditionally dispensable chromosomes in pathogenic *Fusarium* species (Ma et al. 2010) have influenced the perception of genomic evolution in fungi. They have shown that HGT can provide a faster way to gain the ability to colonize new hosts and expand metabolic capabilities than vertical inheritance.

Existing studies of fungal HGT have mostly focused on the Dikarya fungi (Richards et al. 2011; Marcet-Houben and Gabaldón 2010), while HGT in the early diverging fungi (EDF, non-Dikarya fungi) is relatively understudied. Although no comprehensive characterization of HGT in EDF has been presented so far, it is already known that it does occur and may have a profound impact on the recipient organism. Recent reports show that EDF experienced several documented waves of transfers, like the acquisition of secondary metabolite clusters in *Basidiobolus* (Tabima et al. 2020), *Mortierella* (Wurlitzer et al. 2021a) and *Rhizophagus* (Venice et al. 2020), or number of enzymes transferred to *Neocallimastigomycota* involved in the metabolism of nucleotides (Alexander et al. 2016) and carbohydrates (Murphy et al. 2019). A more complex scenario was reported for *Rhizophagus irregularis* that acquired genes both from the host plant and its endosymbiotic bacterium (Li et al. 2018). Catalase genes of bacterial origin, which presumably increase ROS resistance, were found in the amphibian pathogen *Batrachochytrium dendrobatidis* (Hansberg, Salas-Lizana, and Domínguez 2012).

In this work, we study the patterns of gene exchange between fungi and non-fungal organisms on a representative set of 44 EDF proteomes ranging from *Chytridiomycota* to *Mucoromycota*. We have designed a custom phylogenetic-based pipeline (see **Fig 1**. and https://github.com/mciach/HGTin44EDF) to obtain a reliable sample of xenologs, focusing on limiting the number of false positive results that distort the observed signal (Irwin et al. 2022) while retaining a sufficient sample size to infer statistical trends. As potential sources of false positives, we have focused on genome contaminations, spurious homology and promiscuous domains, the discrepancy between closest blast hits and phylogenetically nearest neighbors, unreliable alignments, long branches, weak bootstrap supports, uncertain taxonomy and sequence homoplasy. We then analyze the resulting data set of HGTs from a taxonomic, ecological, and molecular perspective. Our results provide a comprehensive characterization of the patterns of HGT across the EDF tree of life.

**Fig 1.**
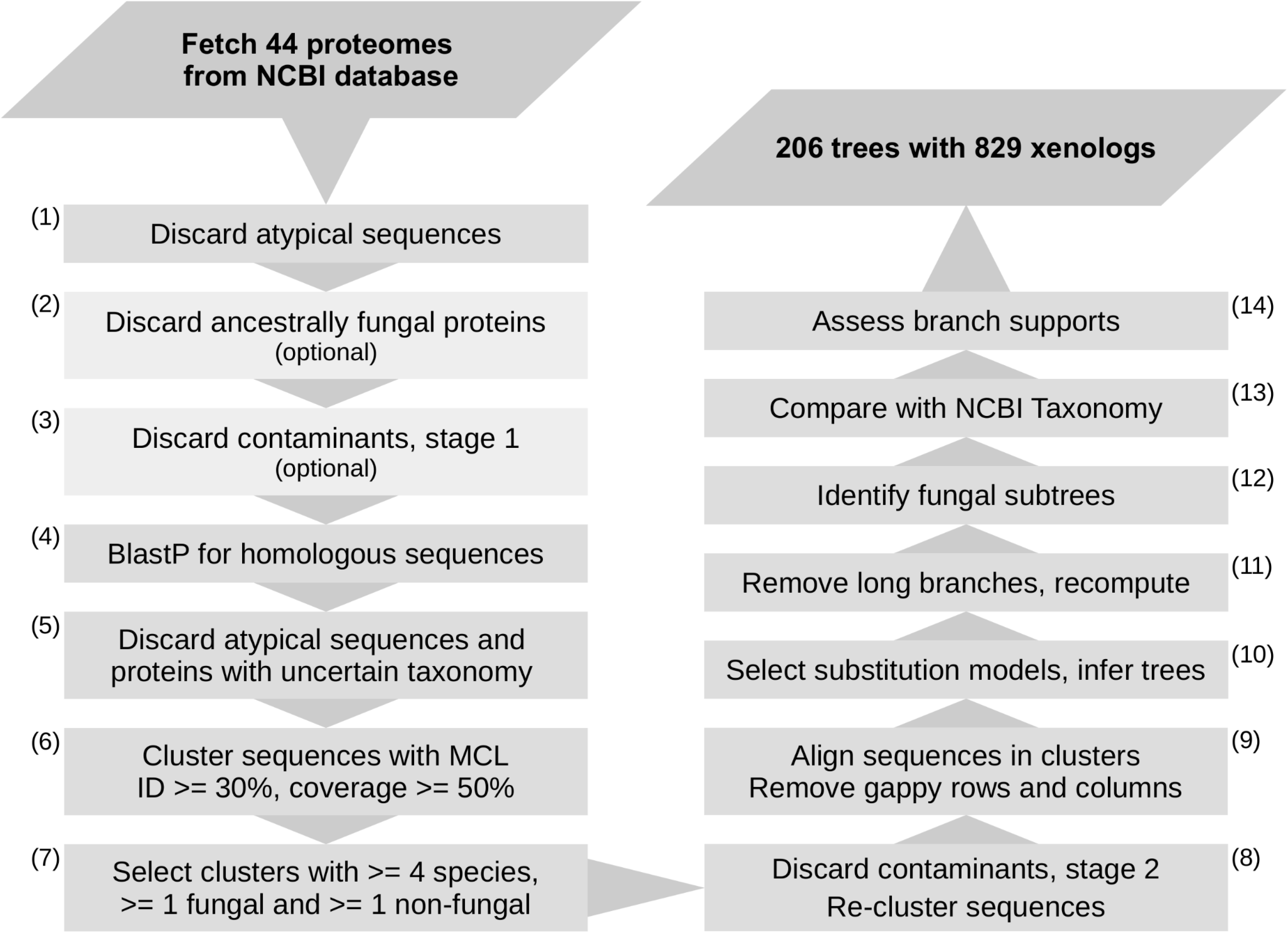
A flow chart of the HGT identification pipeline.

## Methods

### Dataset

Proteomes of 44 early diverging fungi, referred to as the *target proteomes* (Supplementary Table S1), were downloaded from NCBI in April 2018 (NCBI Resource Coordinators 2017). The proteomes jointly contained 606201 proteins. Fungal ecological tagging was obtained from Fungal Traits (Põlme et al. 2020). Taxonomical annotations were obtained from the NCBI taxonomy database, with manual fine-tuning of taxonomic names and ranks when needed according to (Voigt et al. 2021a). Since the NCBI Taxonomy tree contains many polytomies, we take a conservative approach and report HGTs only if they are supported by all possible binarizations. Genomic features, including intron count, were obtained from the GFF files of the genome assemblies. Assembly completeness was assessed with BUSCO v5. BUSCO scores (Manni et al. 2021) and N50 values are given in Supplementary Table S1.

### HGT identification

Our HGT identification approach is visualized in Fig 1. A more detailed description of the rationale and algorithms used in selected steps are available in the Supplementary Material. First, low-complexity regions in all target proteins were masked using ncbi-seg with relaxed settings (window length 12, trigger complexity 1.8, extension complexity 2.0) (John C. Wootton and Federhen 1993). Target proteins with less than 30 non-masked amino acids, more than 2000 characters, or more than 50% masking were discarded as prone to produce false-positive hits (step (1) in Fig. 1). This is because long proteins often have a complex multi domain structure, with domain shuffling and dynamic domain loss/gain, which impacts both alignment quality and phylogeny inference (Stolzer et al. 2015). This procedure removed 5949 out of 606201 proteins.

Note that, although it increases the reliability of prediction, it alsocomes at a cost of missing some of the well documented transfers of long proteins, such as the transfer of NRPS to Mortierella species (Wurlitzer et al. 2021b). However, since less than 1% proteins are removed overall, it is unlikely to have a noticeable influence on the statistical patterns. Target protein homology within the 44 target proteomes was evaluated using the blastp suite with an e-value threshold 1e-05 (NCBI Resource Coordinators 2017). Target proteins with homologs in multiple fungal families were discarded as likely ancestrally fungal (step (2) in Fig. 1). Contigs without any protein coding gene with detectable homology to the other 44 target fungal proteomes were discarded as likely genome contaminants (first-stage contaminant filtering; step (3) in Fig. 1).

Sequences of proteins remaining after step (3) were blasted against a local copy of the Non-Redundant NCBI database (last updated December 01, 2020), using blastp suite with an e-value threshold 1e-04 and maximum of 500 target sequences. Detected homologs were masked using ncbi-seg (default settings). Proteins with less than 30 non-masked amino acids, more than 2000 characters, or more than 75% masking were discarded (step (5) in Fig. 1). The taxonomic lineage of each homolog was inspected using the NCBI Taxonomy database. Only homologs from bacteria, archaea and eukarya were retained in order to discard viral and artificial sequences due to their uncertain location in the species tree (step (5) in Fig. 1). Proteins with ‘unclassified’, ‘uncultured’, ‘environmental’, or ‘incertae sedis’ NCBI Taxonomy keywords anywhere in their lineage were discarded because their placement on the species tree is undefined (step (5) in Fig. 1). An important exception here is that proteins from the 44 target fungal proteomes analyzed in this work were not subjected to this filtering step, because some of the 44 fungi belong to incertae sedis families, and therefore applying this filter would fully discard their proteomes.

All remaining proteins were clustered with MCL (step (6) in Fig. 1). First, an all-vs-all BlastP was performed, with an increased word size to increase specificity and speed up computations (-word_size 6 -threshold 21 -evalue 1e-06). Only HSP matches with identity at least 30% and covering at least 50% of both the query and the subject proteins were retained. This step was essential to avoid cluster merging caused by long, multi-domain proteins and by small but highly conserved regions. The resulting protein homology graph was clustered with MCL (inflation parameter = 2). The results were filtered by discarding clusters without target proteins, with less than 4 different species, with species composition being more than 60% fungal, and clusters with fungal sequences from more than a single phylum. These clusters were assumed to be either unlikely to contain HGTs, ancestrally fungal, or exhibiting evolutionary patterns too complex for an automated analysis (step (7) in Fig. 1).

For each target protein remaining after step (7), we have inspected the contig of the protein’s coding gene in order to discard potentially contaminating sequences (step (8) of Fig. 1). This step can be just as well performed at any previous stage of the analysis; however, performing it after the initial filtering of sequence clusters greatly decreases its computational cost. For each contig containing a gene encoding a target protein, 10 proteins were selected (or, if the contig contained less than 10 proteins, all of its proteins). The selected proteins were blasted against the NR database. Contigs with no detected protein coding region with a fungal first blast hit were discarded as likely contaminants. The proteins remaining after this filtering step were clustered again with MCL using the same strategy. The re-clustering was performed because removing large numbers of nodes from a homology graph can alter the structure of clusters.

The clusters resulting from step (8) were aligned using mafft 3.7 (Nakamura et al. 2018) with E-INS-i strategy, recommended for the alignment of general sequences due to having fewer assumptions than other strategies (--genafpair --maxiterate 5000). Alignments were then filtered by discarding constant columns and columns with more than 80% gaps, and next by discarding sequences with less than 20 non-gap characters or with more than 75% gaps (step (9) in Fig. 1).

Next, maximum likelihood (ML) trees were inferred using IqTree (Minh et al. 2020) with automated model selection (step (10) in Fig. 1). Branch supports were assessed with UFBoot, 2000 replicates. Leaves were discarded if no fungal leaf was closer than 3.33 substitutions per site (corresponding to an average of 15% sequence identity (Grishin 1995)). This step was performed in order to reduce the complexity of the trees and focus on the neighborhoods of fungal sequences. Next, long branches (both terminal and internal ones) were removed iteratively until all branches were no longer than 1.598 substitutions per site (corresponding to an average of 30% sequence identity; step (11) in Fig. 1). This step was performed in order to limit the influence of long branches on the tree topology. Resulting trees were retained if they contained proteins from at least one target and three non-fungal species. See the Supplementary Material for a detailed description of the branch-cutting algorithm.

For alltrees remaining after step (11), their sequences were re-aligned and the alignments were filtered using the same strategy. Next, ML trees were recomputed using the same strategy. This step was performed because removing sequences can alter the topologies, branch lengths, and branch supports of ML trees.

For each tree, a corresponding species tree was downloaded from a local copy of the NCBI Taxonomy database. For each gene tree/species tree pair, all fungal subtrees were identified (step (12) in Fig. 1). Next, the species composition of the neighboring non-fungal subtrees was compared between the trees (step (13) in Fig. 1; see Fig. 2 for a visual description and the Supplementary Material for a detailed description of this step). Fungal subtrees which had different locations in the gene tree and the species tree were considered displaced and therefore indicative of HGT. Otherwise, if the location of a fungal subtreeagreed in both trees, it was deemed as a possible vertical inheritance or sequence homoplasy and therefore not sufficient evidence to infer a horizontal gene transfer. In the case of polytomous species trees, we have inspected all their possible binarizations and inferred HGT only if it was supported by all binarizations.

**Fig. 2.**
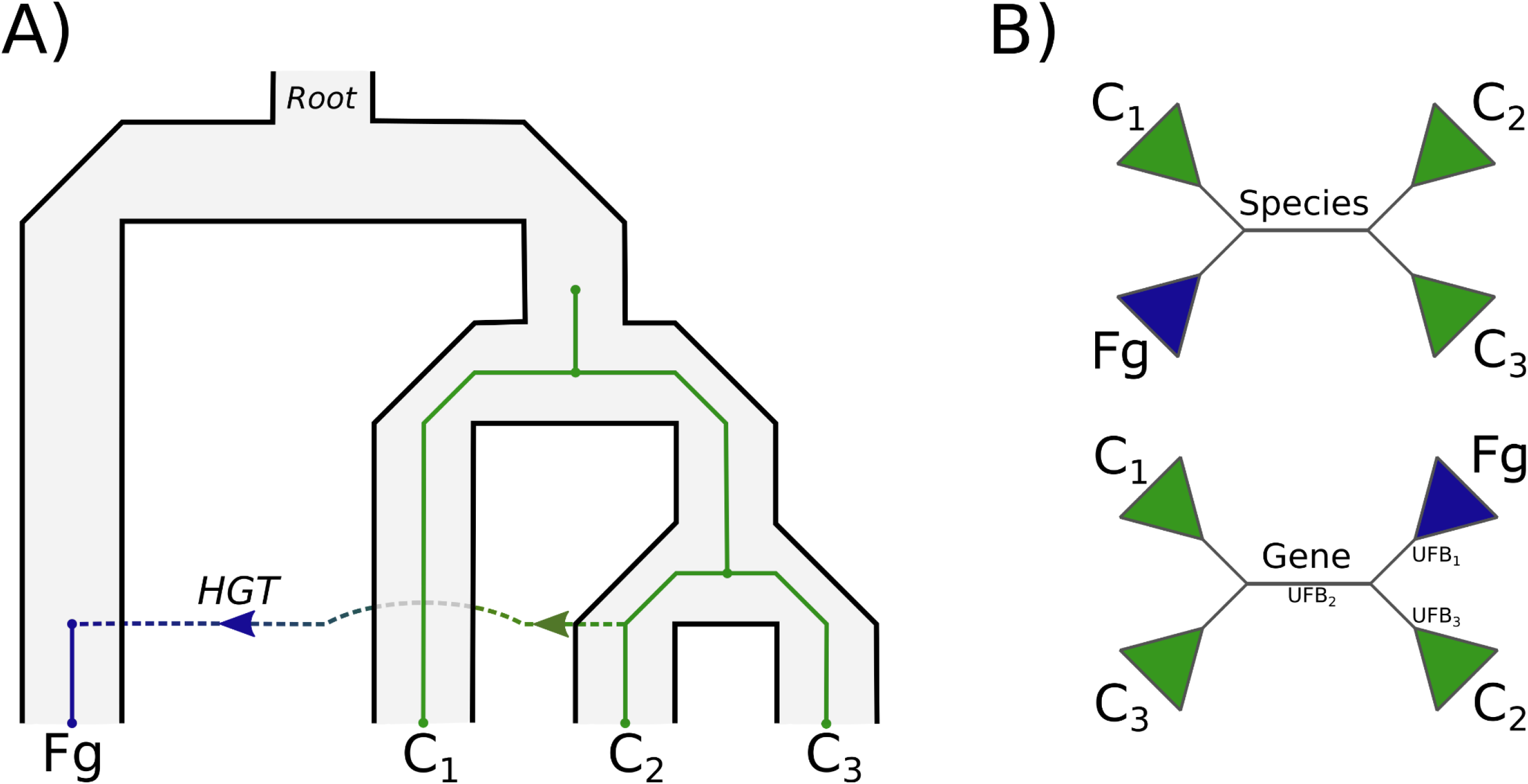
An example of a displaced fungal subtree indicating a horizontal gene transfer. A) A species tree with an embedded gene tree, with a single horizontal gene transfer. B) The same species tree (top) and gene tree (bottom) in their unrooted versions. The fungal subtree {Fg} branches with a {C1} and a {C2, C3} subtree in the species tree, and with a {C2} and a {C1, C3} subtree in the gene tree, indicating a different location of the {Fg} subtree in both trees caused by a horizontal gene transfer into the fungal clade. The correct location of the fungal subtree {Fg} in the gene tree is given by the branch adjacent to the {C1} subtree. The subtree {C2} can be inferred as the donor group because the species LCA of {C1, C3} is the same as the species LCA of the whole neighborhood {C1, C2, C3} (see the Supplementary Methods for a detailed discussion). Trees with an average of the three Ultrafast Bootstrap supports UFB_1_, UFB_2_ and UFB_3_ below 90% are discarded in this study.

To further limit the number of false positive results, we discarded gene trees in which fungal subtrees could not be repositioned to their correct locations, i.e. gene trees with no branch giving the same location (measured in terms of species composition of neighboring subtrees) as in the species tree (see **Fig. 2**). In gene trees without such a branch, any location of the fungal subtree would be artifactually indicative of an HGT. This approach allows for other transfers and evolutionary events within the neighboring subtrees, but discards trees with transfers between the neighboring subtrees or gene tree inference errors as too complex for an automated analysis and prone to return false positive results.

For each fungal subtree, an average of its UFBoot support value and the support of the two neighboring subtrees was computed (step (14) in Fig. 1; see also **Fig. 2**). Candidate HGT fungal subtrees for which the average support of the three branches was lower than 90% were discarded as weakly supported HGTs. For the remaining subtrees, we have estimated the origin of the transfer by inspecting the last common ancestors (LCAs) of species in its two neighboring subtrees and the joint LCA of both neighboring subtrees, ignoring fungal species. If one of the neighbors had the same LCA as the whole neighborhood, we assumed that the other neighbor was the putative donor (see **Fig. 2** and the **Supplementary Material** for a more detailed discussion). This effectively selects the taxonomically smaller neighbor as the donor. Note that this procedure only approximates the donor group in case of additional evolutionary events or gene tree inference errors influencing the neighboring subtrees’ LCAs. In case when both neighbors had different LCAs than their joint LCA, we assumed that the fungal subtrees had an undetermined origin and discarded it from the analysis. Such situations can arise in complex evolutionary scenarios with multiple transfers.

### Analysis of xenolog features

Xenolog protein products were mapped against PFAM (El-Gebali et al. 2019) using pfamscan.pl with default parameters (only hits with significance=1 taken), mapped against KEGG using kofamscan (Aramaki et al. 2020), scanned for signal peptides with TargetP (Armenteros et al. 2019) and cellular localization with WolfPsort II (Horton et al. 2007). For the latter two methods, the result for a given fungal subtree was assumed to be consistent if it was the same for at least 75% of the clade’s proteins. Low-complexity regions were predicted with seg (J. C. Wootton and Federhen 1996) and intrinsically disordered regions with IUPred3 (Erdős, Pajkos, and Dosztányi 2021). A species tree of the 44 target fungi was inferred from whole proteomes with OMA2 based on OMA groups of orthologs (Altenhoff et al. 2019). To assess the evolutionary history of xenologs after transfer, we have reconciled the fungal subtrees (rooted at the transfer acceptor node) with the OMA2 tree in the Duplication-Loss (DL) model using the ete3 Python package (Page and Charleston 1997). Statistical and phylogenetic analyses were performed with the R statistical package (https://www.R-project.org/) and the Python 3 programming language using the Jupyter Notebook (Beg et al. 2021) and numpy (Harris et al. 2020), scipy (Virtanen et al. 2020) and ete3 (Huerta-Cepas, Serra, and Bork 2016) libraries. Plots were created using the matplotlib (Bisong 2019) library. Early diverging fungi taxonomy in this manuscript follows Voigt et al. with Mucoromycota, Mortierellomycota and Glomeromycota treated as separate phyla (Voigt et al. 2021).

## Results

We have detected 829 xenologs in the 44 target fungi with well-supported phylogenetic evidence for a horizontal gene transfer. The xenologs were nested within 226 subtrees in 206 phylogenetic trees. Including other fungal species, the 226 subtrees contained 1208 fungal xenologs. For over 50% of trees, iqTree selected the LG model as the best-fitting one, followed by WAG, VT and JTT models, which jointly accounted for 90% of clusters (**Supplementary Fig. S1**). Nuclear models accounted for 95% of clusters, followed by chloroplast (3%), mitochondrial (1.3%), viral (0.4%).

Additional 283 trees with 326 fungal subtrees containing 254 target xenologs and 1857 xenologs from other fungal species had weakly supported or incomplete phylogenetic evidence for HGT and were not used for the statistical analysis of patterns of HGT. The causes for lack of support are summarized per clade in **Supplementary Fig. S1.** Finally, 109 trees contained 123 fungal subtrees with 282 target proteins which had a congruent location on the gene tree, indicative of a simple vertical inheritance, sequence homoplasy or HGT from fungi rather than HGT into fungi. Note that these scenarios are not mutually exclusive in gene trees, as a single tree may contain multiple subtrees with strong and weak support for HGT as well as subtrees in congruent locations.

The list of all identified xenologs, including ones discarded from analysis due to insufficient support, is available in the Supplementary Material. In the remaining part of this work, unless explicitly stated otherwise, by fungal subtrees we understand subtrees with fungal proteins in gene trees with well-supported phylogenetic evidence for HGT, and by xenologs we understand proteins from target fungi in those subtrees. In this work, we refrain from using the term “transfer event” commonly used in bioinformatic literature, because it suggests one biological event of gene exchange which may result in transferring multiple genes, and its connection with protein families and gene tree subtrees is difficult to establish.

### Genome contamination is an order of magnitude more common than HGT in the analyzed proteomes

In this work, we have adopted a two-stage blastp-based contamination filtering. In the first step, we used a coarse filter in which we discarded contigs without any protein coding gene with a protein product homologous to the other 44 target fungal proteomes. Since this filter is insufficient to detect contaminants with distant fungal homologs, we then refined the filtering by discarding contigs in which we did not find a protein with a fungal top blast hit (excluding the protein’s host).

The first step removed 19578 proteins (**Supplementary Fig. S2**). This constitutes 3% of the 44 proteomes jointly, and 2% of each proteome on average. The number of removed proteins varied highly between proteomes, from 1 in *Absidia repens* NRRL 1336 and *Spizellomyces punctatus* DAOM BR117 through 203 in *Basidiobolus meristosporus* CBS 931.73 up to 5426 in *Rhizophagus irregularis* A1 and *Rhizophagus irregularis* DAOM 197198w. The extensive contamination of the *Rhizophagus irregularis* A1 proteome is in part likely caused by an associated *Lysinibacillus* bacterium.

The second, refined step removed additional 292 proteins. Here, the removed numbers varied to a lesser extent, with the largest being 107 for *Batrachochytrium dendrobatidis* JAM81; notably, the first stage discarded only 36 proteins in this fungus. This was followed by 57 and 53 proteins further removed from *Rhizophagus irregularis* DAOM 197198w and *Rhizophagus irregularis* A1. In all the remaining fungi, the second stage removed less than 10 proteins, suggesting that the initial coarse filter was mostly sufficient to discard contaminants in most fungi. The numbers of sequences discarded in both stages were highly correlated across the 44 proteomes (⍴=0.95), indicating that genome quality is the deciding factor.

The vertical evolution of paralogs after integration into the host’s genome can be described with a Yule-Simons distribution. Over half of the fungal subtrees had only a single leaf, i.e. contained a single fungal protein (**Fig. 2**). If the numbers of xenologs evolved under a neutral birth-and-death model of gene duplications and losses with similar rates of those events across fungi, the distribution of subtree size would follow a geometric distribution (Nee, May, and Harvey 1994). However, this seems not to be the case, and the chi-square test of goodness of fit performed on subtrees with 1 to 12 proteins rejects the hypothesis of a geometric distribution (p<1e-06; ML estimated geometric distribution parameter 0.47). Instead, the distribution of clade sizes appears to closely follow the Yule-Simon distribution with parameter alpha=1 (chi-square test p=0.40; **Fig. 2**), originally derived to describe the number of species in a randomly selected genus (E., F., and Udny Yule 1925; Steel and McKenzie 2001). This distribution differs qualitatively from the geometric one by having a heavy tail, i.e. noticeable occurrences of extremely large values. This gives a theoretical prediction of massive post-transfer duplications independent of transposon bursts, and hints at a considerable variance in the post-transfer duplication rates between HGT events. In fact, the number of post-transfer duplications also seems to follow a Yule-Simons distribution (chi-square p=0.33 for the distribution parameter alpha=2; Fig. 2).

Characteristically for heavy-tailed distributions, while 80% of fungal subtrees in our data set contained up to three proteins, 1.3% contained more than 50, with the largest one containing 219 *Mucoromycota (sensu Mucoromycotina in classification proposed by (Spatafora et al. 2016))* xenologs, with26 in the target fungi. Since the 219 homologs are contained within a single subtree, they seem to be an example of a massive duplication of foreign genetic material during its vertical evolution after incorporation in the host genome. The target proteins from this subtree contain only an uncharacterized domain (DUF4379); other proteins from this cluster are involved in DNA repair and recombination (K03746).

### The number of xenologs is highly divergent across fungi

The number of xenologs in different proteomes was also heavily tailed, with 5 fungal species accounting for over 80% of all detected xenologs (**Supplementary Fig. S3**). These included the gut-dwelling *Neocallimastigomycota* (with xenologs constituting more than 1% of their proteomes), the chytrid *Rhizoclosmatium globosum* (0.5%), and the soil saprotrophic Linnemania *elongata* (0.5%). These groups were followed by the plant-symbiotic *Glomeromycota* and parasitic *Chytridiomycota* (with xenologs forming less than 0.2% of their proteomes). We did not observe any statistically significant correlation between the proteome size and the number of xenologs (⍴=0.19, p=0.21).

The heavy-tailed distribution of post-transfer duplications only partially explains these numbers. Each of the aforementioned fungi seems to have its own evolutionary and ecological mechanism for acquiring large numbers of xenologs. In chytrid species, each strain was the recipient of multiple transfers, while the xenologs in *Neocallimastigomycota* were mostly transferred to the ancestor of the whole group. Both groups were involved in multiple HGT events, while the large number of xenologs in Linnemania *elongata* was caused by only a few transfers and extensive post-transfer duplications.

The average proportion of xenologs in proteomes was 0.12%. This figure is within the range of 0.00% to 0.38% (with an identical per-genome average) reported previously for Dikarya (Marcet-Houben and Gabaldón 2010). We remark, however, that describing heavily tailed distributions with averages is highly inaccurate and may be misleading. The average proportion of xenologs may indeed be 0.12%, but, according to our data, a typical fungus is expected to have none or few at most. Furthermore, this proportion is expected to vary highly between reports due to a particular sensitivity of heavy-tailed distributions to dataset selection.

### Transfer rates are highly divergent across the fungal tree of life and do not always reflect the numbers of xenologs in the proteome

To analyze the distribution of HGTs across the fungi, we have generated an OMA2 phylogenomic tree of the 44 EDFs and labeled its nodes with the numbers of the corresponding fungal subtrees in our HGT data set (**Fig. 2**). Note that this procedure only approximates the depth of the transfer in the tree due to possible post-transfer gene losses or incomplete taxon sampling (making the transfer appear more recent), or inter-fungal transfers after gene acquisition (making the transfer appear more ancient).

The resulting distribution of HGTs across the fungi was highly uneven. There was a clear pattern dividing ancestrally aquatic taxa grouped in *Chytridiomycota* and *Neocallimastigomycota* (living in the digestive tract of terrestrial herbivores) with higher HGT rates compared to typical terrestrial fungi. The mainly saprotrophic *Mucoromycota*, despite the highest sampling in our dataset, shows the lowest number of HGT events per genome, just as the animal-related *Kickxellomycota* fungi. *Mortierellomycota*, despite Linnemania *elongata* having one of the highest numbers of xenologs in our data set, appears to have rather low transfer rates, just as the closely related *Glomeromycota*. As opposed to ancestrally aquatic fungi, post-transfer duplication seems to be the major force driving the number of xenologs in these fungi.

### Transfer rates reflect physical proximity

Diverse bacteria were the most common donors of xenologs in our data set, including *Firmicutes* (most prominently *Clostridiaceae, Lachnospiraceae, Eubacteriaceae, Oscillospiraceae* and *Bacilli*), *Proteobacteria* (*Gammaproteobacteria, Burkholderiales*) and *Actinobacteria*. These were followed by eukaryotic Oomycota, Metazoa and Viridiplantae, and only a few transfers from Archaea (**Fig. 2**). In agreement with previous results (Alexandre et al. 2021), in *Neocallimastigomycota* we have detected genes from a myriad of diverse bacteria, including *Bacteroidetes, Bacilli (Firmicutes), Slackia (Actinomycetota)* and *Aeromonas (Proteobacteria)*. The *Firmicutes* bacteria were a particularly characteristic donor for these fungi, with numerous genes obtained from the anaerobic *Clostridia*, which, according to previous reports, may have enabled them to adapt to the gut environment (Murphy et al. 2019).

For *Mortierellomycota* fungi, where *Burkholderia*-related endobacteria (BRE) are common, *Burkholderiales* stand out as one of the main sources of xenologs. Putative gene donors include *Mycoavidus cysteinexigens* (OAQ22128.1), an endohyphal symbiont of *Linnemannia elongata* (J. Uehling et al. 2017), and *Mycetohabitans rhizoxinica* species (OAQ29846.1), which were previously described as endohyphal symbionts of *Rhizopus microsporus*. The second main source of transfers to *Mortierellomycota* were the soil-inhabiting *Actinomycetia*. This bacterial group was recently reported as the most prevalently associated with fungal isolates (Robinson et al. 2021).

In the water-living *Chytridiomycota*, most xenologs come from loosely associated bacteria such as *Alphaproteobacteria, Gammaproteobacteria* and *Actinomycetia*. In addition, we have identified several transfers from Oomycota, including a tRNA (N6-threonylcarbamoyladenosine(37)-N6)-methyltransferase TrmO (ORY41869.1) and tandem tetratricopeptide repeat proteins (e.g. ORY27572.1). TrmOs are widely present in most domains of life but were apparently lost in Fungi. The transfer of the Oomycota gene seems to have compensated for this loss. The repeat proteins are predicted to form a protein-binding surface, but their function remains unknown.

### *Neocallimastigomycota received* multiple independent transfers of homologous proteins

Out of 206 trees in our data set, 187 contained a single fungal subtree, 18 contained two, and one contained three. One of the trees with two subtrees indicated two independent transfers of a protein with trypsin domain into Conidiobolus coronatus, but the average bootstrap support of all internal branches was equal to only 70. All the remaining trees with two or three fungal subtrees contained only Neocallimastigomycota species. In 7 trees with two fungal subtrees and in the single tree with three, all the subtrees contained different sets of species, indicating independent transfers of similar proteins after the divergence of *Neocallimastigomycota*. An example is an independent acquisition of proteins with a solute binding domain *SBP_bac_3* by *Neocallimastix californiae* G1 from *Oscillospiraceae* (ORY20588.1) and by *Anaeromyces robustus* from *Methanobacteriaceae* (ORX65467.1). The fact that these fungi occupy a similar ecological niche could predispose them to independently receive similar genes through HGT, resulting in a kind of a convergent evolution. On the other hand, in four trees, the two subtrees contained identical species, indicating two independent transfers to the ancestral lineage of *Neocallimastigomycota*. An example is a repeated acquisition of a bacterial cellulase by the ancestor of *Neocallimastigomycota*, once from *Bacilli* (e.g. *Neocallimastix californiae* G1 ORY56492.1) and once from *Clostridia* (e.g. *Neocallimastix californiae* G1 ORY21818.1). This result stands in agreement with a recently reported major HGT event in the ancestor of *Neocallimastigomycota* that might have driven its evolution towards a lifestyle as gut-dwelling fungus of herbivores (Murphy et al. 2019). In the remaining 7 trees, the species composition overlapped partially between the two subtrees, indicating a repeated ancestral transfer followed by gene losses.

### Low-complexity sequences are rarely transferred

The distributions of the proportions of protein sequences covered by low-complexity regions, as well as intrinsically unstructured regions, have lighter tails for xenologs than for the proteome background (**Fig. 5**). A comparison of the average low-complexity proportions of fungal xenologs to the putative donor group showed no visible trend (**Supplementary Fig. S4**). This suggests that extensive low-complexity or unstructured regions tend to inhibit transfers (or retainment of the foreign genetic material), rather than being caused by them. Note, however, that it may be an artifact caused by impaired homology detection between low-complexity regions (Jarnot et al. 2022). The distribution of xenolog sequence length follows the background up to around 1800 amino acids and drops sharply afterwards. This seems to be an artifact of our method, which puts a threshold on xenolog sequence length at 2000 aa. Individual cases of transfers of much longer sequences are documented, for instance, secondary metabolic clusters in *Basidiobolus* and multiple Dikarya (Tabima et al. 2020; Wisecaver, Slot, and Rokas 2014). As a consequence, we cannot conclude whether the distribution of sequence length of xenologs follows the protein background or not and whether sequence length poses a barrier for HGT in fungi. A comparison of the average sequence length of xenologs to the putative donor group showed no visible trend of shortening or lengthening of sequences post-transfer, but revealed differences of up to 400 amino acids in both ways (Supplementary Fig. S4).

### Xenologs are secreted more often than typical fungal proteins, but determining their precise subcellular location may be challenging

TargetP 2.0 detected a signal peptide in 33% of identified xenologs, compared to 7.7% of the proteome background. This stands in agreement with the osmotrophic lifestyle of fungi, their massive secretome, and the accessory function of many xenologs. Consistently with the bacterial origin of many xenologs, only 9 xenologs had mitochondrial transfer peptides. We did not observe biologically meaningful differences in the numbers of transmembrane helices predicted by TMHMM, although a chi-square test weakly rejects the hypothesis of identical distributions in xenologs and the background (p=0.03, ddof=0; assuming xenolog observations as observed, background as expected).

The subcellular localization predicted by WolfPsort was consistent for 77% of target protein subtrees, compared to 95% for TargetP (**Fig. 5**, **Supplementary Fig. S5**). Compared to the proteome background, xenologs were predicted to localize more often in the extracellular space, the endoplasmic reticulum, and the cytoplasm, and less often in the nucleus and the mitochondria. Most xenologs localized in the extracellular space, closely followed by the nucleus. In contrast to TargetP, WolfPSort predicted 90 proteins to localize in the mitochondria. The unexpectedly high number of mitochondrial and nuclear predictions may suggest that WolfPsort is biased towards these locations. To verify this hypothesis, we have generated a decoy dataset by randomly permuting each sequence from the 44 proteomes. The predicted distribution of locations was very similar for the proteome background and the decoy dataset, with the nucleus as the most common location, and the decoy proteins attained similar prediction scores (**Supplementary Fig. S5**). This was likely caused by the fact that WolfPSort is based on the *k*-Nearest Neighbours classification algorithm, which tends to reflect the distribution of the training dataset, in this case, fungal proteomes. The prediction may be further complicated by the fact that xenolog sequences have a non-fungal origin.

### Xenologs typically contain few introns but can acquire many

Over 50% of xenologs did not contain any intron, and almost 90% contained at most two, compared to respectively 25% and 59% in the background (**Fig. 5**). This result is in line with recommended HGT verification measures (Jaramillo, Sukno, and Thon 2015). However, we have also detected 16 xenologs, mostly of eukaryotic origin, with 6 or more introns (**Fig. 5**). In particular, the *Glomeromycota* fungus *Rhizophagus irregularis* DAOM 197198w and the chytrid fungi *Gonapodya prolifera*, *Batrachochytrium dendrobatidis* JAM81 and, to some extent, *Rhizoclosmatium globosum* were prone to have intron-rich xenologs from both unicellular eukaryotic and putative prokaryotic donors. Although there are some chytrid species referred to as *intron-rich chytrids* (van de Vossenberg et al. 2019), neither *G. prolifera* nor *R. globosum* have been described as such in their genomic papers (Y. Chang et al. 2015; Mondo et al. 2017). The preference for the intron structure of xenologs seemed to some extent correlated taxonomically: while the xenologs in *Chytridiomycota* tended to contain multiple introns, the ones in *Neocallimastigomycota* tended to contain few if any (Fig. 5 C). The number of introns closely followed a geometric distribution in the fungal proteome background, but not in the set of xenologs. In particular, the distribution in xenologs was zero-inflated, consistent with the bacterial origin of many xenologs. However, many bacteria-derived xenologs contained multiple introns, suggesting that they acquire introns after the transfer, consistent with previous reports (Lage et al. 2013). This seemed to be the case in an *R. globosum* xenolog with twelve introns (ORY15655.1), derived from a deltaproteobacterium *Labilithrix luteola* magnesium transporter. This protein had three post-transfer paralogs (i.e. R. globosum proteins nested in the same fungal subtree), out of which one contained only one intron, one contained five, and one contained six (respectively ORY29294.1, ORY29292.1 and ORY15656.1). A similar scenario seems to have shaped the story of arginine decarboxylase biosynthesis gene, which was transferred from an intronless gene of an anaerobic deltaproteobacterium *Anaeromyxobacter dehalogenans* (locus_tag A2cp1_2068) and acquired nine introns after transfer to *R. globosum* (KXS13513.1). In addition to the intronization of bacteria-derived xenologs, we have also observed an apparent intron gain in an E3 ubiquitin-protein ligase gene with 8 introns, transferred from some early diverged metazoans to *Glomeromycota* (EXX59914.1). The closest metazoan homologs from *Acropora millepora* (LOC114972538) and *Orbicella faveolata* (LOC110047255) have only three or four introns.

To test the hypothesis of intron acquisition in bacteria-derived xenologs, we have compared the number of introns to the branch distance to the root of the corresponding fungal subtree (i.e. the post-transfer number of substitutions per site). A linear model fitted without an intercept estimated 0.6 introns per unit branch length (p < 0.001). However, the overall picture is more complicated, as some xenologs quickly acquired multiple introns, while many others had none even after a long time.

### Gene transfer seems to frequently involve acquiring and losing protein domains

Compared to the proteome background, fungal sequences obtained *via* HGT were more likely to map to one or more distinct PFAM domains (**Fig. 5**). The average number of domains was 0.85 for the background and 1.11 for the xenologs.

However, one hundred fungal xenologs had domains which had no clan-level siblings in non-fungal sequences from the same gene tree, and were therefore apparently acquired after the incorporation of the foreign genetic material in the host genome. Most xenologs acquired a single domain, while 10 acquired two distinct domains. Except for one xenolog in Linnemania *elongata* AG 77 and five xenologs in chytrids, all the post-transfer domain acquisitions happened in *Neocallimastigomycota*. As therefore expected, the by far most commonly added domain was the carbohydrate-binding module CBM_10 (PF02013), a part of the dockerin cellulosome typical of anaerobic fungi, acquired in 86 xenologs. Consistent with the function of CBM_10, this domain was fused preferentially to xenologs containing domains associated with carbohydrate metabolism, such as the bacterial glycosyl hydrolase (PF02011) in 31 xenologs. Some exceptions to this rule involved 13 xenologs with a bacterial CotH kinase domain, the main invasin involved in mucormycosis which binds to glucose-regulated protein 78 (GRP78) on nasal epithelial cells (K. B. Nguyen et al. 2016; Alqarihi et al. 2020). The next most commonly acquired domain was the ricin B lectin domain (PF14200) acquired in nine *Neocallimastigomycota* xenologs.

Some domains were seemingly lost during transfer, with 89 xenologs not containing domains which were present in at least 20% of sequences in both neighboring gene tree subtrees. An example was a bacteria-derived cellulase precursor ORX60185.1 in *Piromyces finnis*, which was missing an N-terminal fragment of its bacterial homologs with a cellulose binding domain CBM_2 and a Bacterial Ig domain Big_7. On the other hand, this protein contained a bacterial glycoside hydrolase domain Glyco_hydro_48 and was post-transfer fused with a cellulose binding domain CBM_10. Effectively, a bacterial carbohydrate-binding domain was replaced with a fungal one in this protein.

Overall, out of 829 xenologs detected in our target fungi, 170 had a novel or a lost domain, and 48 had both (such as the aforementioned cellulase precursor in *P. finnis*). In some cases, this resulted in conflicting phylogenetic signals. The *R. globosum* ORY15655.1 (one of the aforementioned intronized bacteria-derived xenologs) contained a xenologous N-terminal region of 530 aa of bacterial origin with a glycoside hydrolase domain (InterPro *Glycoside_hydrolase_SF*) and an apparently ancestrally fungal C-terminal region of 180 aa with a mainly eukaryotic magnesium transporter NIPA domain Mg_trans_NIPA. We note that this apparent gene fusion may have simply been a gene calling error merging two neighboring genes (frequently encountered in genomes without transcriptomic evidence and submitted as drafts), and requires further proof for confirmation. Regardless of the actual source of the apparently ancestrally fungal region, the presence of conflicting signals in this protein’s sequence severely impacts the topology and support of the phylogenetic tree (**Supplementary Fig. S6**).

Notably, the 12 introns of the R. globosum ORY15655.1 xenolog were located in both the ancestrally fungal and the bacteria-derived region. The three aforementioned putative post-transfer paralogs of this protein with fewer introns (ORY29294.1, ORY29292.1 and ORY15656.1) lack the apparently ancestral C-terminal region (**Fig. 4**). Whether the presence of an ancestrally fungal genetic material has facilitated the intronization of ORY15655.1 is an open question. Interestingly, the three paralogs also lacked any PFAM or InterPro domains, including the bacteria-derived InterPro glycoside hydrolase domain.

**Fig. 3.**
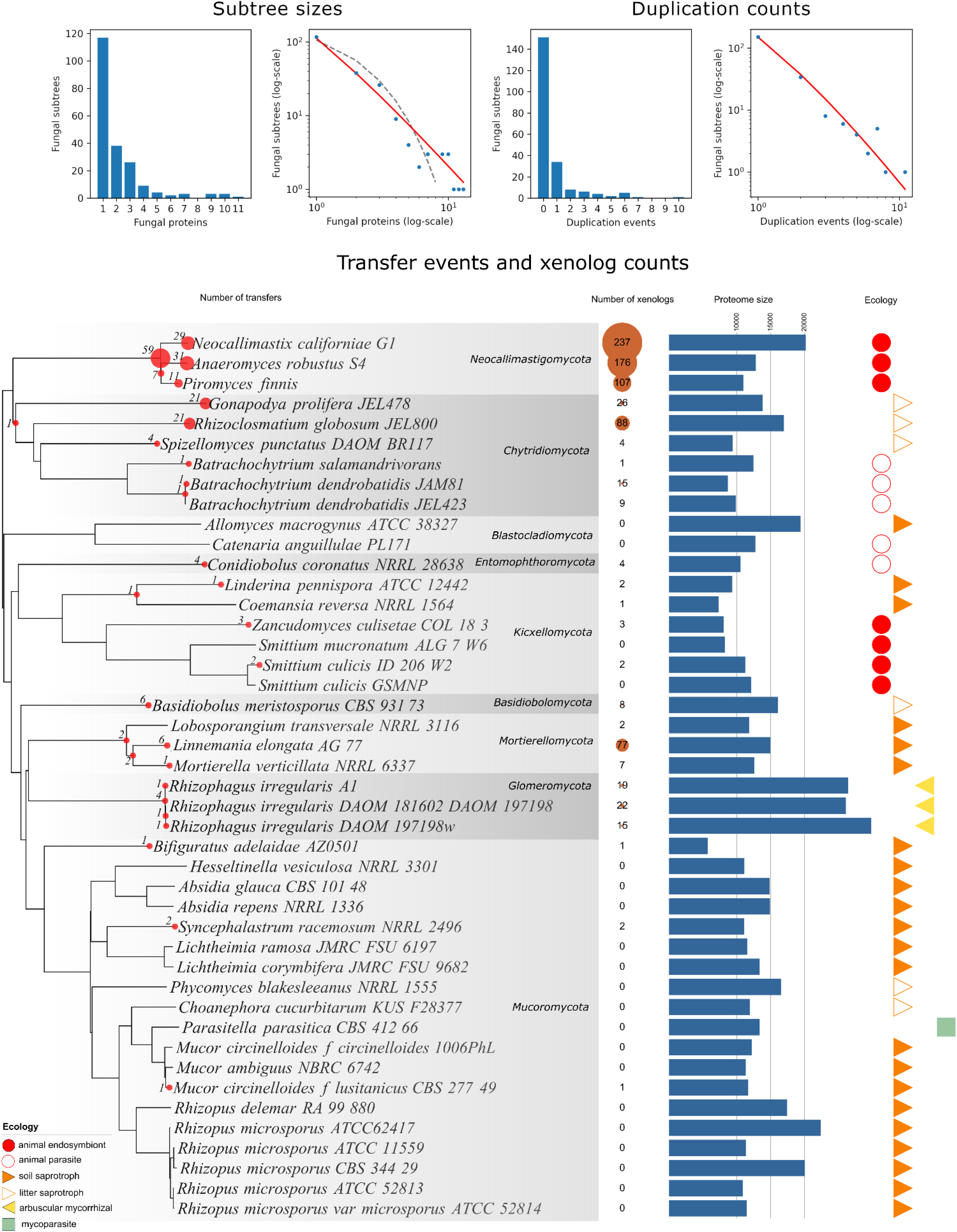
The rates of HGT and post-transfer duplications among the 44 EDF. A) The fungal subtree sizes and the numbers of post-transfer duplications among the target fungi. The red lines on the log-log plots correspond to the Yule-Simons distributions (parameter alpha=1 for subtree sizes, alpha=2 for duplications), the dashed gray line corresponds to a geometric distribution with parameter 0.47. B) The OMA2 phylogenetic tree of fungi annotated with the numbers of subtrees of xenologs (brown bubble) and of transfers (red dots on tree, their size is proportional to the number of events). Proteom size and ecology of fungus is given on the right.Taxonomic names in B) are based on NCBI Taxonomy with manual fine-tuning according Voigt et al. (2021a). For the position of the outgroup and Dikarya see Voigt and coauthors. (Voigt et al. 2021b).

**Fig. 4.**
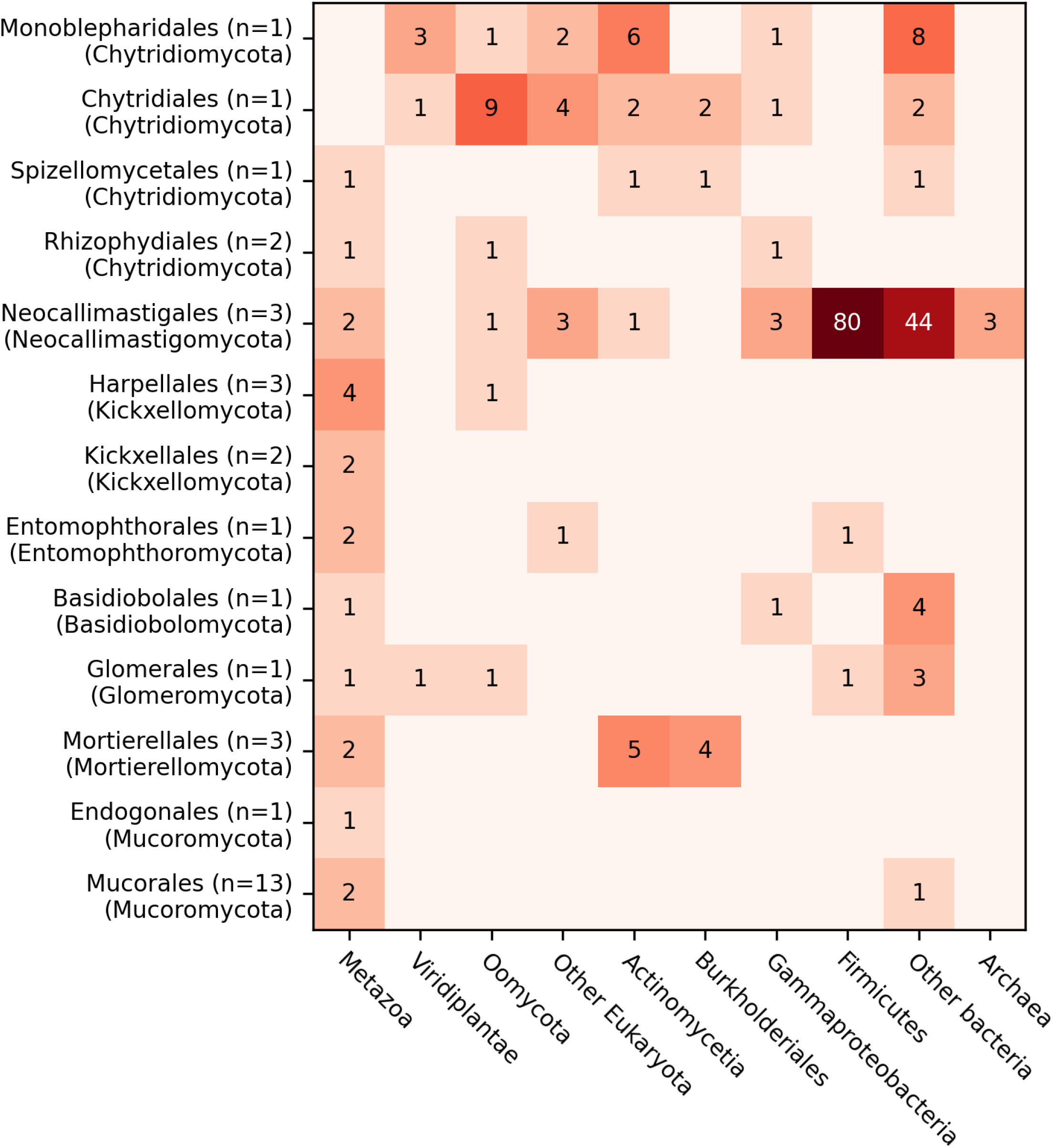
Numbers of subtrees per acceptor-donor pair.

**Fig. 5:**
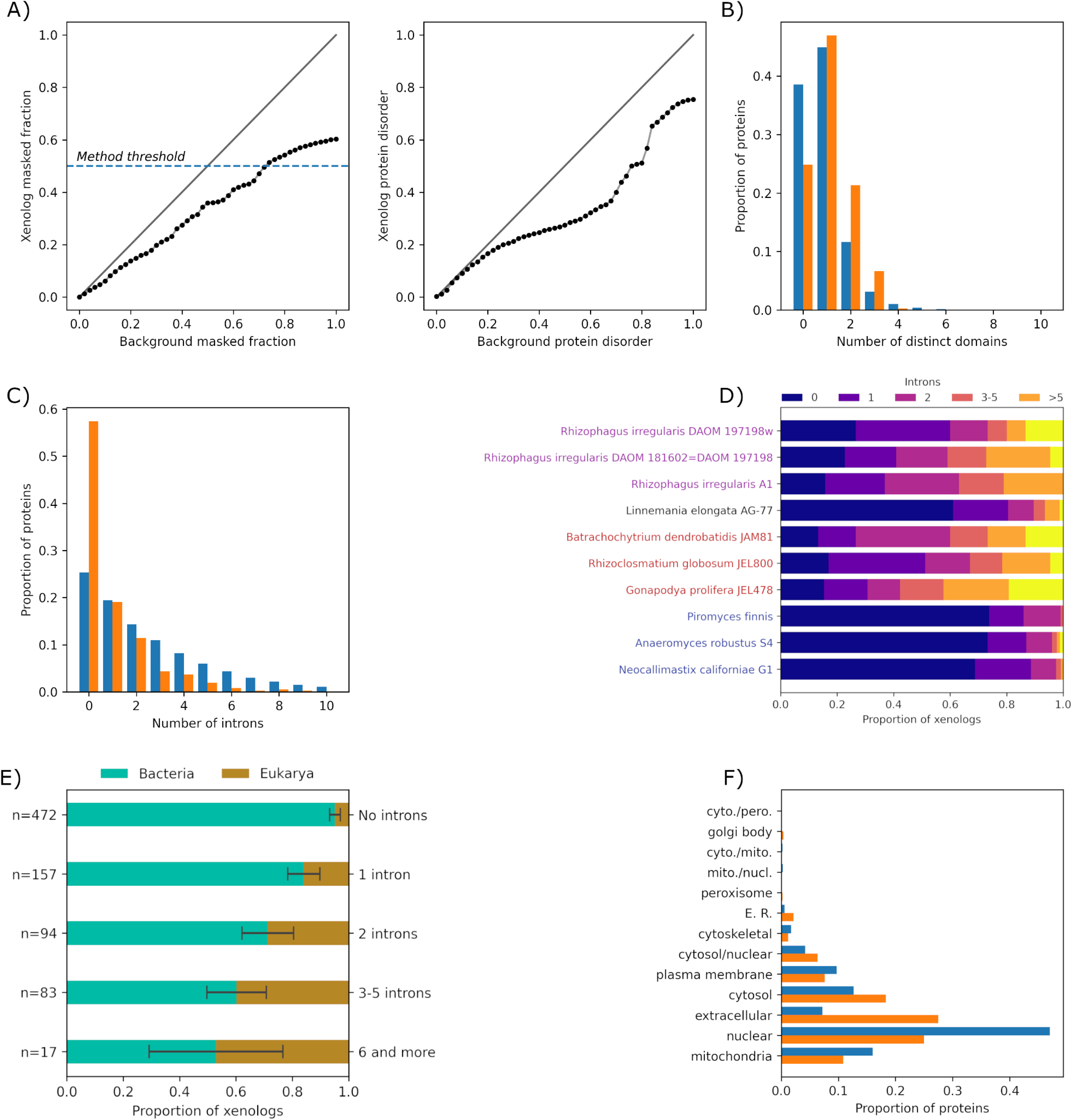
Molecular characteristics of xenolog protein products (orange) and background proteins (blue). A) Quantile plots comparing the xenolog and background sequence low-complexity fraction and intrinsically unstructured fraction. B) The numbers of identified PFAM domains. C) The numbers of introns. D) A detailed distribution of intron counts in xenologs of selected fungi, showing taxonomy-correlated preferences. Glomeromycota species highlighted in magenta, Chytrid species in red, Neocallimastigomycota species in blue. E) The number of introns and their dependence on the kingdom of the putative donor. F) The subcellular localizations of xenologs and the background proteins.

**Fig. 6:**
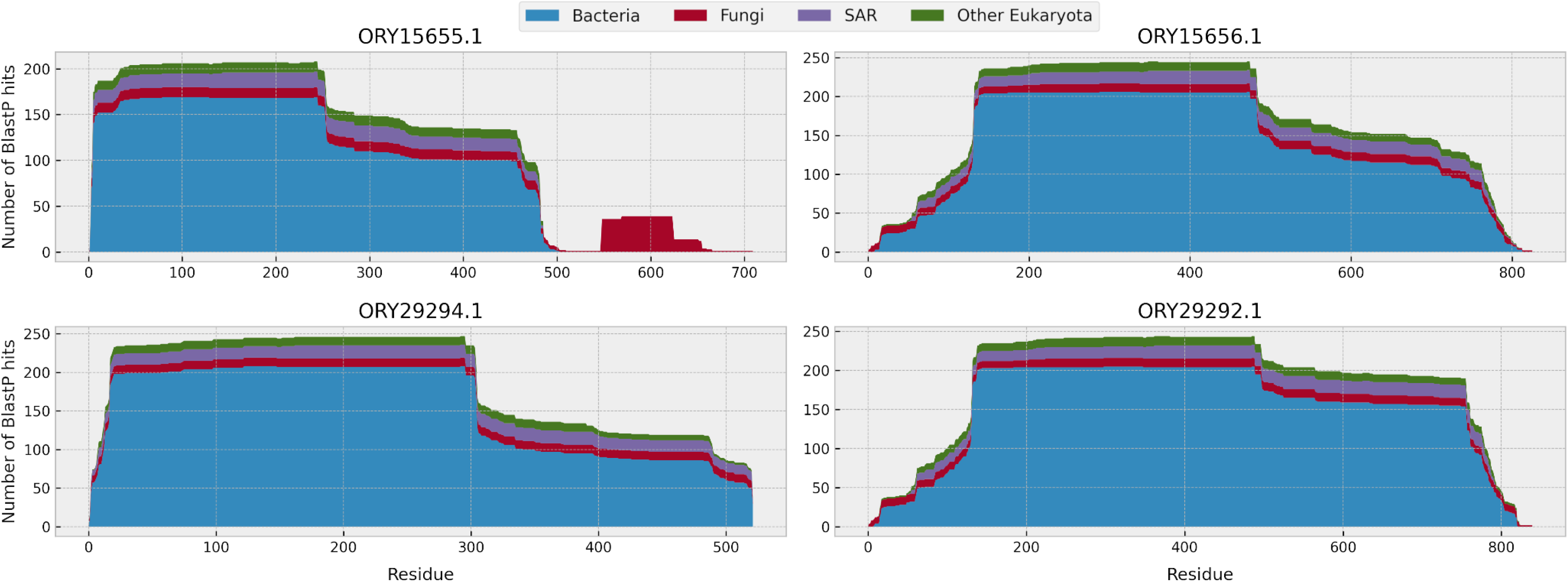
A post-transfer fusion of a fungal domain with a xenolog. Plots visualize the taxonomic composition of the first 250 BlastP hits per amino acid of the query protein. The ORY15655.1 protein contains a distinct C-terminal region which is likely ancestrally fungal and contains a magnesium transporter domain (residues 548-623). The post-transfer paralogs of this protein (ORY15656.1, ORY29294.1, ORY29292.1) have a similar taxonomic distribution of their BlastP hits at each amino acid but lack the domain.

### Metabolic domains constitute the majority of xenologs

To analyze HGT-derived protein domains of fungal xenologs, we have selected domains which were present in at least half of fungal proteins in a given subtree and in at least 20% of proteins in one of the neighboring subtrees. Domains of unknown function (DUFs/UPFs) constituted a numerous functional class with 16 diverse Pfam families found in 60 xenologs. Among domains with known functions we have found numerous transfers of genes involved in metabolism, in particular coding alpha/beta hydrolases, acetyltransferases and esterases. Many carbohydrate active enzymes, such as glycohydrolases and glycosyltransferases, were transferred to *Neocallimastigomycota (as reported previously by Murphy and coauthors (Murphy et al. (2019))*. Peptidases were underrepresented in the dataset (peptidase xenologs n=24) compared to the carbohydrate active enzymes and carbohydrate binding modules (n=63), but were transferred to diverse recipients. They included trypsins transferred from *Phytohabitans flavus* to *Spizellomyces punctatus* (KNC97026.1) and from *Araneus ventricosus* to *Conidiobolus coronatus* (KXN68345.1) and a membrane-bound dipeptidase, metalloproteinase M19, transferred from *Micromonosporaceae* to *G. prolifera* JEL478 (KXS08983.1). Trypsin expansions can be linked to enhanced degradation of protein-rich substrates such as insects, while a M19 peptidase is a part of an *Aspergillus* spp. gliotoxin biosynthesis cluster. However, the *Aspergillus* spp. M19 peptidase is likely ancestrally fungal and distant to the *G. prolifera* M19 peptidase.

The dominance of carbohydrate active enzyme encoding genes over protein metabolism genes in our data set can be linked to the high number of transfers to *Neocallimastigomycota*. These fungi also acquired genes with a myriad of specificities connected to secondary metabolism and transport, extending their nutritional capabilities. Condensation domain (PF00668) containing candidate NRPS secondary metabolite clusters were transferred from *Eubacteriales* to several *Neocallimastigomycota* (e.g. *Anaeromyces robustus* ORX81431.1). General transporters from MFS_1 and ABC families were also acquired by these fungi (eg. ORY49205.1).

We have detected a few transfers (n=7) of mobile elements, including a transfer of a PiggyBac element from *Phytophthora* to *Smittium culicis* (OMJ27075.1) and of a retrotransposon from *Globisporangium splendens* to *Rhizoclosmatium globosum* (ORY50945.1). However, most of the potential transposon xenologs were likely excluded from our data set because of the usage of protein sequences instead of genomic sequences for HGT inference, as well as our thresholds for branch support values,branch lengths, and taxonomic distributions. We have also identified several phage proteins without clear function in fungi, eg. a phage tail lysozyme, PF18013, in *Piromyces finnis* protein ORX49608.1.

We have detected transfers of several genes involved in resistance, including a bacterial putative betalactamase resistance gene transferred from to *G. prolifera* JEL478 (KXS12969.1). *Mortierellomycota* acquired NOD-like receptors with NACHT-PPR-WD40 architecture from *Mycoavidus cysteinexigens* endosymbionts. NOD-like receptors are suspected to play a role in Dikarya immunity (Jessie Uehling, Deveau, and Paoletti 2017; Dyrka et al. 2014). Interestingly, all of the *Mortierellomycota* immunity proteins in our dataset seem to be of a bacterial origin or at least shared with the endosymbiont. *Mycoavidus* homologs have ∼50-45% of sequence identity and are the best reciprocal blast hits. It is an open question whether these proteins have a role in the symbiotic interaction.

On the other side, we have detected transfers of several putative toxins. Several *Neocallimastigomycota* representatives gained a bacterial ricin-type sugar binding modules (18 xenologs in a single subtree; e.g. ORX83988.1, ORY77981.1). Most of those domains were associated with bacterial glycoside hydrolase domains and nearly all were post-transfer fused with the carbohydrate binding CBM_10 domain. Additional nine Neocallimastigomycota xenologs in another gene tree appear to have gained the bacterial ricin-type sugar binding module post-transfer, suggesting a proliferation of a xenologous domain across the host’s proteome (e.g. ORX84139.1, ORY73345.1). Similar proteins were recently documented in Mucor acting as toxins (mucoricin) (Soliman et al. 2021). Among other examples, a sea anemone pore-forming toxin (PF06369) was transferred from *Selaginella moellendorffii* to *R. irregularis* and was present in all three strains analyzed in this study (e.g. PKC66727.1).

### Putative sequence homoplasy has a similar taxonomic distribution to HGTs

Annotation of the OMA2 tree of target fungi with the numbers of non-displaced fungal subtrees in our data set (indicative of vertical inheritance or sequence homoplasy) revealed a striking similarity to taxonomic patterns of HGTs (**Supplementary Fig. S7**). The correlation between the numbers of displaced and non-displaced subtrees mapped to the nodes of the OMA2 tree was 0.80 (p < 1e-12). Note that, due to taxonomic filters applied in our workflow, the species composition in those trees was at least 60% non-fungal (in terms of both the number of proteins and the number of distinct taxa). This finding suggested that, for fungal proteins with a predominantly non-fungal homology, a HGT may be a more likely hypothesis than vertical inheritance with multiple gene losses or sequence homoplasy even in the absence of tree incongruence. Although homoplasy has been reported to occur frequently on the scale of amino acids in Eukaryota (Rokas and Carroll 2008), it seems unlikely on the scale of proteins. Therefore, the 282 target proteins discarded in this work solely due to lack of tree incongruence are likely to represent additional cases of HGT. They include e.g. *Mortierellomycota* fatty acid desaturases similar to delta-12 fatty acid desaturases of bacterial origin, likely acquired from Deltaproteobacteria (KFH67875.1). The representatives of this fungal group are known to produce long polyunsaturated fatty acids (L. Chang et al. 2021).

## Discussion

Our aim in this work was to provide a statistical characterization of the patterns of interkingdom HGT across diverse EDF. In this kind of a high-throughput study, there are multiple factors that can inflate the false positive rate of HGT detection (see the **Supplementary Methods** for a detailed discussion), including, but not limited to:

- Relying on sequence similarity instead of phylogenetic trees (Koski and Brian Golding 2001)
- Excessively long, multidomain proteins, sometimes originating from ORF prediction errors (Poptsova and Gogarten 2010),
- Sequences without a well-defined location in the species tree (e.g. viral or environmental),
- Low-quality alignments and a wrong choice of the trimming software (Tan et al. 2015),
- Automatic rooting of gene trees (Lamarca, Mello, and Schrago 2022; Wade et al. 2020),
- Low-quality gene trees with low support values and long branches (Rodríguez-Ezpeleta et al. 2007),
- Using tree reconciliation algorithms with improper parameters (Libeskind-Hadas et al. 2014).

In order to avoid those pitfalls, we have applied a series of filters and developed a custom-made algorithm for the detection of gene trees with displaced fungal clades. Notably, our method allows us to use unrooted gene trees to detect transfers and approximately identify their putative donors and does not require the specification of any parameters. Our data and results should provide a reliable source of fungal xenologs and their properties which can further be used to benchmark new HGT identification methods (**Supplementary Table S1**). In particular, statistical properties such as the Yule-Simons distribution of the numbers of xenologs per transfer event could be used to validate future results in a rigorous, statistical framework. This would be a particularly valuable addition to other criteria if a mechanistic explanation of this distribution was provided and verified.

Our results suggest that the false positive risk factors listed above have a greatly varying impact on the quality of the results. Moreover, while there has been much research devoted to some, others seem to be underappreciated in the literature. Phylogenetic tree incongruence is widely considered the gold standard of HGT detection and has been the basis of our approach. However, the exceptionally high correlation between the numbers of congruent and incongruent trees per fungal taxon, combined with the lack of correlation between them and the recipient’s proteome size, suggests that taxonomy-based methods such as the Alien Index may be equally reliable (Rancurel, Legrand, and Danchin 2017; Gladyshev, Meselson, and Arkhipova 2008). The added benefit of phylogeny-based methods is a more precise determination of the putative gene donors. However, it is important to note that, with currently available methods, this precision is always limited by the completeness of the available databases, and the identified putative donors are at best only the closest sequenced relatives of the actual donors.

While tree-based methods seem to offer only a marginal improvement over taxonomy-based heuristics, the filtering of contaminant sequences is of tremendous importance. We have detected nearly 20 000 likely contaminating sequences, 20 times more than the number of detected xenologs. As a consequence, a randomly selected gene with an atypical taxonomy of homologs or incongruent phylogeny is much more likely to be a contaminant than a xenolog. In EDF, genome contamination may occur due to sample contamination or the presence of endohyphal bacteria. The ultimate determination of whether a gene is incorporated into the genome and expressed requires experimental evidence; in bioinformatic analysis, the minimum requirement should be the presence of ancestral genes in the same contig. Heuristic approaches based on sequence identity and GC content may be insufficient to separate xenologs from contaminants, especially in the case of genes transferred from endosymbionts. The picture is further complicated by the fact that fungal endosymbionts are still understudied, and most likely there still are many unknown endohyphal organisms (Robinson et al. 2021).

The state-of-the-art workflows for HGT identification, including the one used in this work, consist of multiple stages, with each stage having multiple possible solutions and parameters. One of the consequences is that results obtained with different methods are seldom comparable. Although the rates and patterns of HGT have been studied in depth in individual lineages of early diverging fungi, such as *Neocallimastigomycota (Wang et al. 2019)*, *Basidiobolus* (Tabima et al. 2020) and *Batrachochytrium (B. Sun et al. 2016)*, each of those studies employed its own methodology. Furthermore, they were exploratory in nature rather than comparative, investigating in depth the potential impact of HGT on a selected lineage. To our knowledge, the work presented in this manuscript is the first one that applies a unified methodology to multiple fungal lineages in order to provide an extensive comparative study across the tree of life of early diverging fungi. This allowed us to show, for example, that *Neocallimastigomycota* are more receptive to HGTs than *Basidiobolus* despite the latter having more HGTs reported in the literature, and more receptive to HGTs than *Linnemania elongata* despite having similar numbers of xenologs in their proteomes.

Our goal to limit the number of false positive results necessarily means that we also lose many true HGT events. Numerous previously reported xenologs failed to pass one or more of our requirements, such as xenologs reported in *R. irregularis* (e.g. EXX72150.1) (Li et al. 2018), which we found to be in a congruent location in the gene tree, numerous xenologs reported in *B. dendrobatidis* such as adenylate cyclase (XP_006679093.1) or chitinase (XP_006682460.1) (B. Sun et al. 2016), which we found to be widespread across chytrids, or octin in *Phycomyces* (XP_018295118.1) (T. A. Nguyen et al. 2018), which we found to be widespread across fungi. The transfer of NRPS to *Mortierella* from *Mycoavidus* (Wurlitzer et al. 2021b) was discarded due to excessive sequence length, although we have identified parallel transfers of other proteins. The transfer of proteins with Gal_Lectin domain to *Neocallimastigomycota* (Wang et al. 2019) was discarded due to lack of homology to non-fungal species. The transfer of an ATPase to *R. irregularis* (ESA12330.1) (Li et al. 2018) was discarded due to low sequence coverage of the homologous region.

On the other hand, our methodology has recovered several previously reported HGTs. Out of numerous xenologs reported in *Neocallimastigomycota* in (Wang et al. 2019), we have identified 10 as strongly supported. Out of 811 xenologs reported in *Basidiobolus* in (Tabima et al. 2020), we have identified 7 as strongly supported, 23 as weakly supported and 6 in a congruent location in the gene tree. As opposed to exploratory studies, focusing on low numbers of uniformly sampled observations is appropriate for comparative statistical analyses. As soon as trends are detectable and testable, adding further observations offers little benefit (provided that sampling of observations is done correctly). Statistical hypothesis testing determines whether the sample size is sufficiently large to distinguish trends from random fluctuations.

After filtering out the contaminants and applying multiple filters to discard false positives and obtain a clear statistical signal, we have shown that one of the most predictable features of EDF HGT is its unpredictability. The number of xenologs in a given proteome is driven by at least two burst-like events: bursts of gene exchange like in *Neocallimastigomycota* and bursts of gene duplications like in Linnemania sp. This agrees with unexpectedly high numbers of xenologs in some metazoan proteomes, like *Rotifera* sp. (Gladyshev, Meselson, and Arkhipova 2008). Nevertheless, we have also identified some general patterns in the properties of protein products of horizontally transferred genes. Xenologs are characterized by relatively short low-complexity and intrinsically unstructured regions. The depletion of low-complexity regions can be related to the so-called complexity hypothesis, which states that xenologs tend to be far from central hubs of interaction networks (Cohen, Gophna, and Pupko 2011), and instead participate in secondary metabolism, drug resistance etc. On the other hand, regions with a skewed amino acid abundance, which constitute 20% of eukaryotic proteins (Mier et al. 2020), are involved in diverse molecular functions usually linked to regulation (Wright and Dyson 2015) and protein-protein interactions (Coletta et al. 2010).

In agreement with the current notions, the xenologs in our dataset tend to contain few introns. We have, however, detected notable exceptions to this rule in chytrids and some *Glomeromycota*. This seems to have resulted from both the acquisition of intron-rich genes and the intronization of xenologs. The distribution of introns seems to be taxonomy-dependent, with certain fungal groups showing a preference for intronless xenologs, while others for intron-rich ones. Bacteria-derived xenologs acquire introns over time, but some are particularly resistant to intronization or not yet ameliorated to the fungal genome. Together with the different distributions of the numbers of introns in xenologs and the proteome background, this suggests that the intronization dynamics is more complex in xenologs than in fungal proteins.

The lack of correlation between the number of xenologs and proteome size implies that HGT has little or no influence on the proteome sizes of EDF. Moreover, large proteomes do not predispose to HGT and the overall amount of genetic material does not seem to be a limiting factor for the incorporation of transferred genes.

The receptiveness for foreign genes varies greatly between fungal lineages, with bacteria being the most common source of xenologs. In agreement with previous reports, we have found that fungi belonging to ancestrally aquatic groups experienced more HGT events than terrestrial lineages. This can be explained by the easier exchange of genetic material in the aquatic ecosystem, the ubiquity of unicellular aquatic life forms, and the possibility of forming biofilms with bacteria (McDaniel et al. 2010; Grüll, Mulligan, and Lang 2018; Abe, Nomura, and Suzuki 2020). Another possible explanation is that terrestrial fungi may have lost some of the HGT predisposing traits or gained defensive mechanisms preventing foreign gene incorporation. Despite the impact of the aquatic environment, the relationship with the host and the necessity to adapt to a new environment seem to have a higher influence on the rate of HGT. Our observations suggest that, while symbiosis or parasitism does not always result in gene exchange, it is a predisposing condition.

Some of the early diverging fungal lineages have associated bacteria, either endohyphal or loosely related to their hyphae, some of which are co-evolving with the fungal host and transmitted vertically (Pawlowska et al. 2018). Those bacteria have been previously pointed to as a likely source of xenologs (Naito, Morton, and Pawlowska 2015), because such close endosymbiotic relations pose a natural opportunity for a large-scale gene exchange, as exemplified by the massive gene transfer from the mitochondrion to the nucleus of eukaryotic organisms (Anselmetti et al. 2021). This pattern was confirmed in this study, with a notable example of *Podila verticillata* acquiring fatty acid desaturases probably from *Deltaproteobacteria* and NOD-like receptors from endohyphal *Burkholderia*-related endosymbiont *Mycoavidus* sp. The latter suggests a parallel evolution of NOD-like receptors and perhaps a NOD-like receptor-based immunity in Dikarya and *Mortierellomycota*. Moreover, NOD-like receptors may be involved in the host-bacteria interaction. *A parallel case of HGT of TIR domain-containing protein coding genes / NOD-like receptors was reported in Gigasporaceae and expressed in response to Gigaspora endosymbiont CaGg (Venice et al. 2020).* Our results also confirm the reports on a massive enzyme encoding gene acquisition by *Neocallimastigomycota* (Murphy et al. 2019) which has likely played a role in their transition to a gut-dwelling mode of life.

Out of 829 xenologs detected in our target fungi, 100 had apparently post-transfer fused domains. This may point to an unexpectedly high frequency of gene fusions occurring during or after the integration of foreign DNA. Cross-kingdom HGT has been reported previously as an important factor in gene fusion (Yanai, Wolf, and Koonin 2002). As the foreign genetic material is incorporated into the host’s genome, it can be located within an existing gene. Alternatively, homologous or non-homologous recombination can occur after the new gene is incorporated, especially if it contains a fragment similar to the host’s genes. While it is unclear which mechanism was responsible for the swapping of the bacterial cellulose binding domain with a fungal one in ORX60185.1, a replacement of a domain which preserves the protein’s function agrees with the current notions about the evolution of domain architectures.

Note that the 100 fusions were detected only based on identified domains; since many sequences lack identified domains, the full extent of post-transfer gene fusion is unknown. Frequent fusions may significantly complicate the studies of HGT. Recombination of transferred and ancestral genetic material may, on one hand, confound the impact of HGT on the host’s ecology and evolution, and on the other hand, the proteins’ phylogenetic signals. While most research to date, including this one, has focused on detecting “whole-protein” xenologs, our results indicate the need to develop domain-level methods of xenology detection and to investigate their results. This agrees with the recent ideas that although terms like orthology, paralogy or xenology are traditionally applied on a level of whole genes or proteins, they should perhaps also be considered on a domain-level basis (Linard et al. 2021; Persson et al. 2019; Sjölander et al. 2011; Forslund et al. 2018). Domain-level HGT detection methods may also be more robust against gene calling errors merging neighboring genes or missing their parts. We also note that sequence regions with independent evolutionary histories are another reason why the first blast hit may not be the phylogenetically closest neighbor, as a small but highly conserved region may have a lower E-value than a large but less conserved one.

In this work, we have focused on xenologs which homology is confined to a single taxonomic family of fungi. This was done in order to obtain clear-cut cases of HGTs in simple gene trees, and therefore to mitigate the false positive rates. However, discarding proteins with homology to more than one fungal family means that our methodology mostly detects relatively recent transfers from distantly related putative donors. In particular, it does not detect transfers between fungal species, such as the massive transfer from a fungal parasite *Parasitella parasitica* to its host *Absidia glauca (Kellner et al. 1993)*, and may also have a limited sensitivity towards transfers of animal genes, many of which have a fungal homolog (like the aforementioned polyubiquitin in *Zancudomyces* (Wang et al. 2016)). As a consequence, while our results confirm the occurrence of cross-kingdom gene transfers between eukaryotes (Schönknecht, Weber, and Lercher 2014; Van Etten and Bhattacharya 2020; Richards et al. 2009), they may significantly underestimate their prevalence.

The list of xenologs reported in this work is far from exhaustive. We have found several additional probable cases of HGT among gene trees discarded due to different reasons in different steps of our pipeline. The ORY16048.1 protein in *Rhizoclosmatium globosum* is a plausible transfer from mosquitoes, which occupy a similar ecological niche as the recipient fungus in their larval states; however, the protein has a low identity to its closest homolog and therefore has been discarded in the long branch cutting stage. One of the possible reasons for low identity is the absence of the actual donor in the NR database; accordingly, progress in genome sequencing efforts is likely to enable the discovery and confirmation of many new HGTs. In conclusion, although much research has already been done, a thorough characterization of the rates and patterns of HGT still requires extensive further work in optimizing HGT detection protocols and algorithms.

## Ethics approval and Consent to participate

Not applicable

## Consent for publication

Not applicable

## Author Contributions

M.C, P.G., J.P. and A.M. designed the study, M.C. and P.G. developed the algorithms, M.C. implemented the software, M.C. and A.M. prepared the dataset and performed genome analyses, M.C. performed the statistical analyses. All authors have interpreted the data and wrote the manuscript.

## Availability of data and materials

Phylogenetic trees of supported and unsupported HGT are deposited under 10.5281/zenodo.7461853, all accessions and assemblies are listed in Supplementary Table S1.

