## Supplementary Material for "The interkingdom horizontal gene transfer in 44 early diverging fungi boosted their metabolic, adaptive and immune capabilities"

**Supplementary Figures** **3**

Supplementary Fig. S1. Left: the numbers of phylogenetic trees with different sequence substitution models fitted by ModelFinder of iqTree. Right: Causes of an incomplete support for the HGT hypothesis in gene trees discarded from the statistical analysis of HGT properties. 3

Supplementary Fig. S2. Numbers of contaminants removed from the 44 EDF proteomes in both stages of contaminant filtering. The first stage discards contigs without any fungal homology, the second stage further discards contigs without any fungal first hit in 10 randomly sampled proteins. 4

Supplementary Fig. S3. The size of the proteome (blue) and the number of detected xenologs (orange) for 44 fungi analyzed in this study. Note the different axis scales for the blue and orange bars. 5

Supplementary Fig. S4. The dynamics of sequence length and low-complexity proportion of proteins after transfer. Left: The difference between the average lengths of the fungal acceptor and the donor group sequences. Right: The difference between the average proportions of sequences masked with ncbi-seg. 6

Supplementary Fig. S5. Additional results of the WolfPSort subcellular location prediction software. A) Location consistency of fungal xenologs in a xenologous family, defined as the fraction of fungal proteins with identical location in a protein cluster. B) The comparison of location prediction for proteome background and decoy proteins generated with random permutations of protein sequences. C) The distribution of WolfPSort prediction scores for the background proteins, decoy proteins, and xenologs. 7

Supplementary Fig. S6. Detailed BlastP results and phylogenetic trees for the protein ORY15655.1, containing a xenologous as well as a short, putatively ancestrally fungal region. The influence of the ancestrally fungal signal is visible as a distortion of the topology of the tree constructed from the whole sequence compared to trees constructed from different regions. The tree constructed from the N-terminal region of 530 aa supports a bacterial origin of the sequence, the tree constructed from the C-terminal region of 180 aa supports a fungal origin of the sequence, while the tree constructed from the full sequence shows a mixed taxonomy. All trees constructed from the first 500 BLASTP hits of respective query sequence using the BLAST web suite (full sequence; residues 1:530; residues 531:710). 8

Supplementary Fig. S7. A comparison of the numbers of incongruent and congruent fungal clades in gene trees with predominantly non-fungal taxonomic composition. Incongruent (displaced) clades are assumed to be a stronger indication of HGT than congruent (non-displaced) clades, as the latter may potentially also be caused by sequence homoplasy and simple vertical evolution. A high correlation of the numbers of both types of clades across the species tree of 44 EDF suggests that HGT is a more likely explanation for the congruent clades, as homoplasy would be expected to produce a uniform distribution over the tree. 9

**Supplementary Methods** **10**

Avoiding false positives in high-throughput HGT studies 10

Controlling long branches in gene trees 12

Detecting horizontal gene transfer in unrooted gene trees 13
References 17

### Supplementary Figures

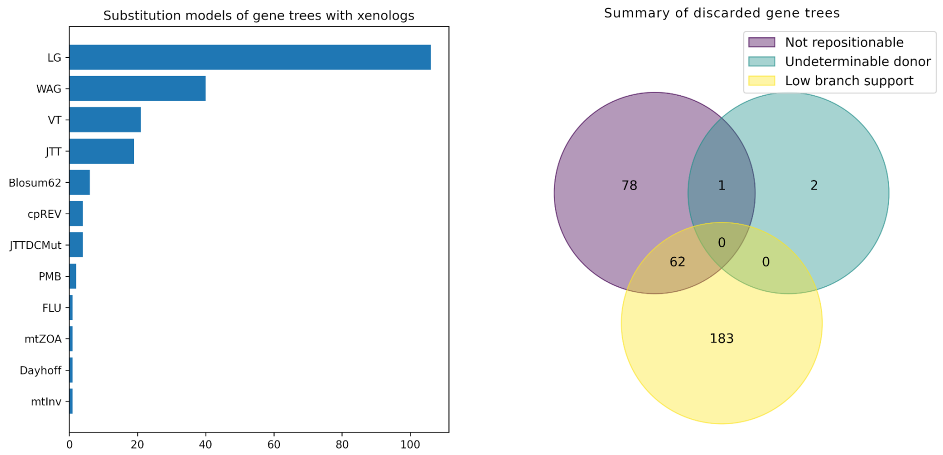


**Supplementary Fig. S1**. Left: the numbers of phylogenetic trees with different sequence substitution models fitted by ModelFinder of iqTree. Right: Causes of an incomplete support for the HGT hypothesis in gene trees discarded from the statistical analysis of HGT properties.

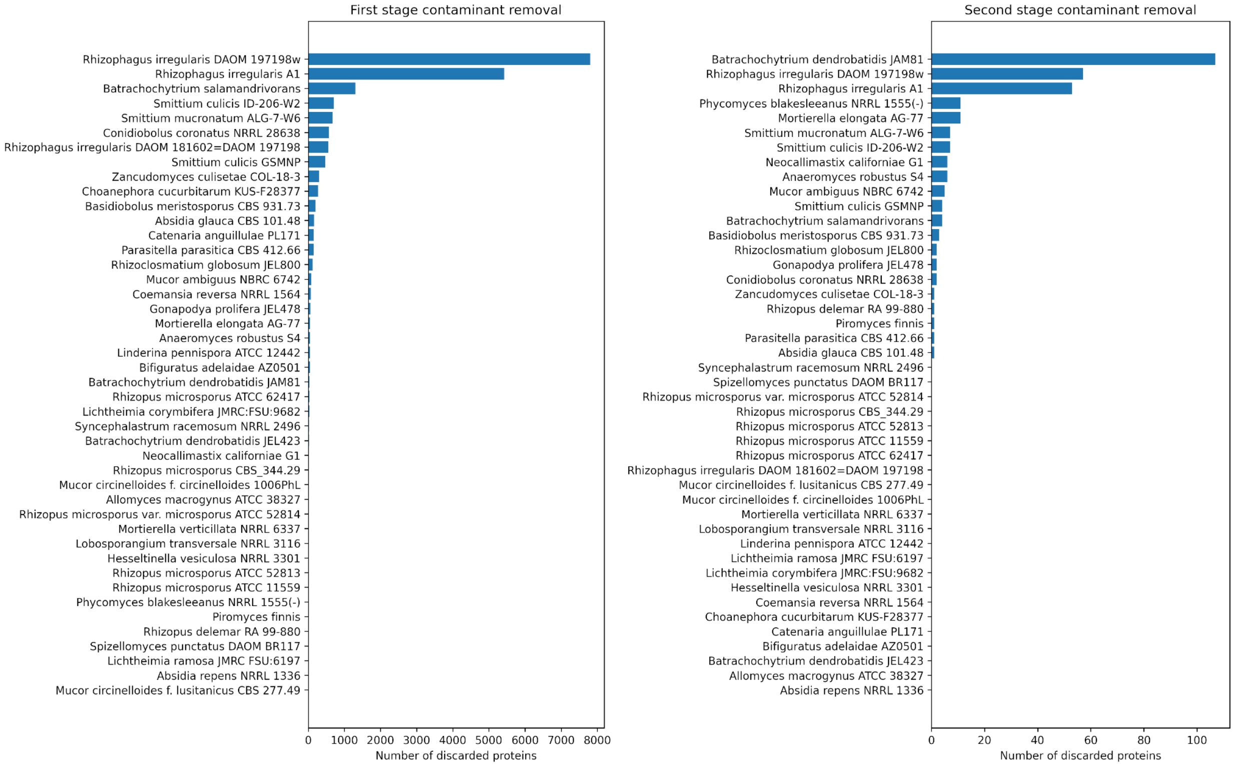


**Supplementary Fig. S2.** Numbers of contaminants removed from the 44 EDF proteomes in both stages of contaminant filtering. The first stage discards contigs without any fungal homology, the second stage further discards contigs without any fungal first hit in 10 randomly sampled proteins.

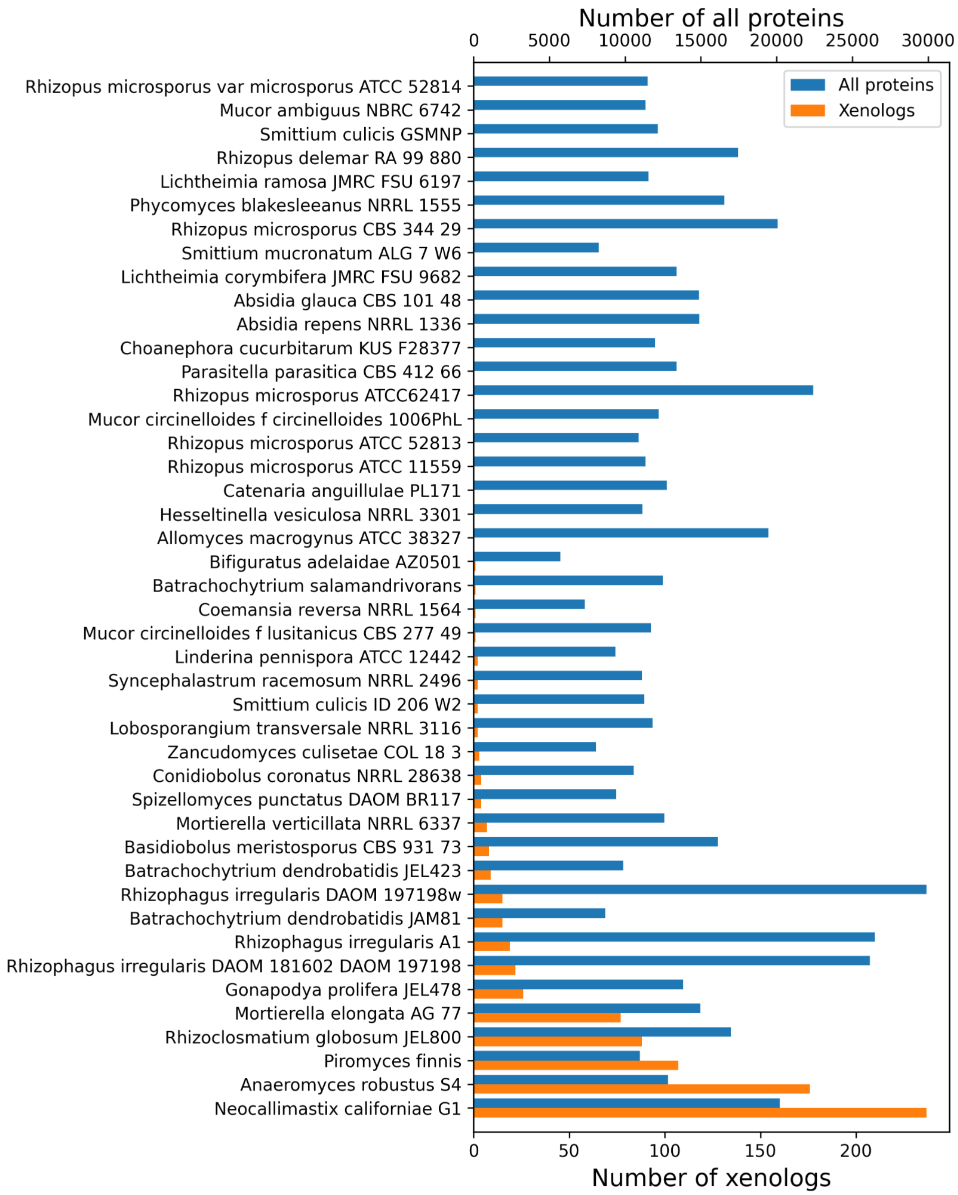


**Supplementary Fig. S3**. The size of the proteome (blue) and the number of detected xenologs (orange) for 44 fungi analyzed in this study. Note the different axis scales for the blue and orange bars.

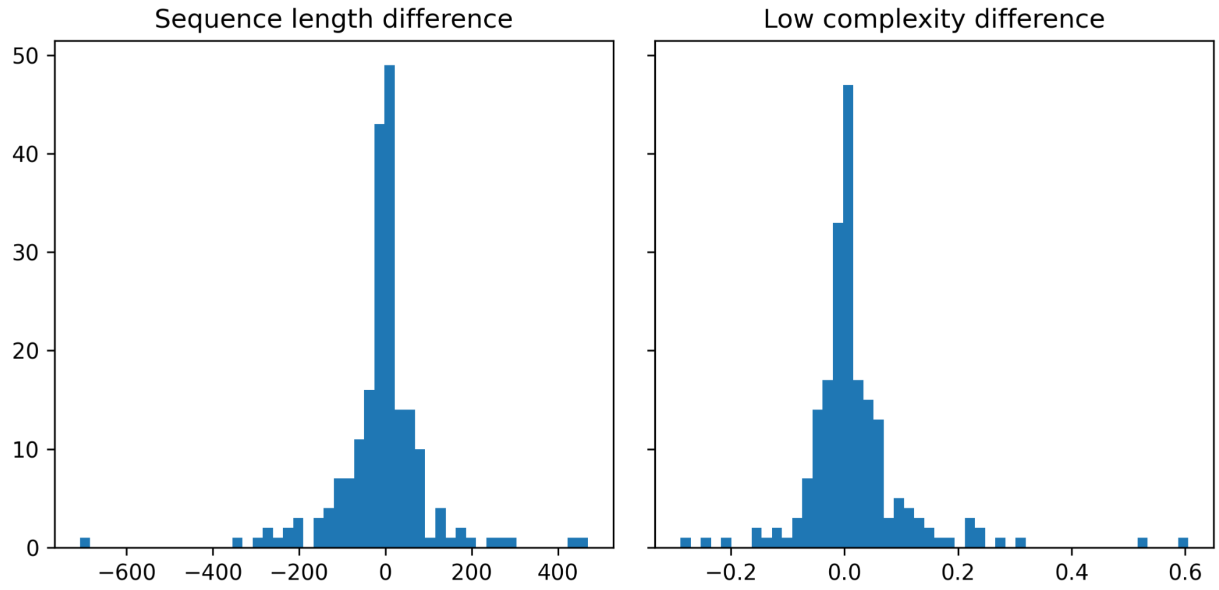


**Supplementary Fig. S4.** The dynamics of sequence length and low-complexity proportion of proteins after transfer. Left: The difference between the average lengths of the fungal acceptor and the donor group sequences. Right: The difference between the average proportions of sequences masked with ncbi-seg.

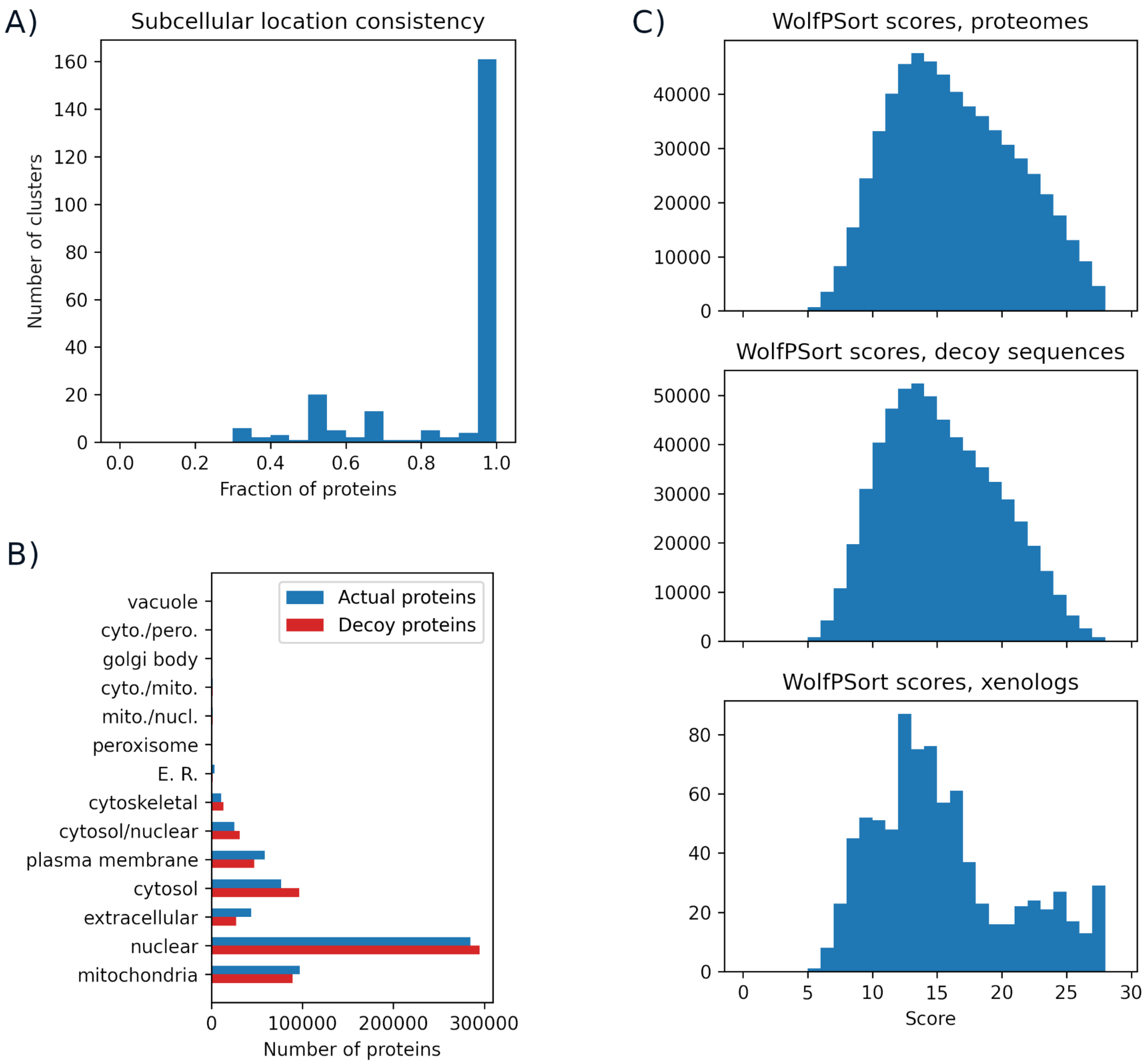


**Supplementary Fig. S5.** Additional results of the WolfPSort subcellular location prediction software. A) Location consistency of fungal xenologs in a xenologous family, defined as the fraction of fungal proteins with identical location in a protein cluster. B) The comparison of location prediction for proteome background and decoy proteins generated with random permutations of protein sequences. C) The distribution of WolfPSort prediction scores for the background proteins, decoy proteins, and xenologs.

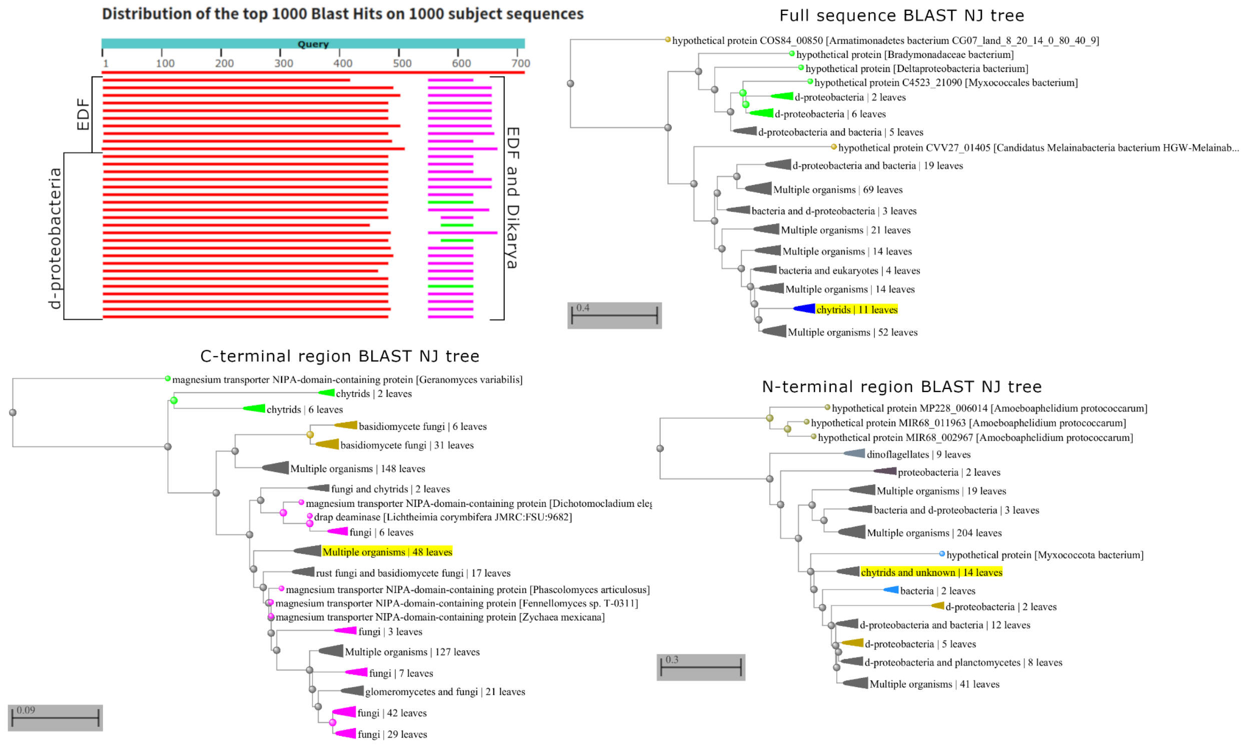


**Supplementary Fig. S6.** Detailed BlastP results and phylogenetic trees for the protein ORY15655.1, containing a xenologous as well as a short, putatively ancestrally fungal region. The influence of the ancestrally fungal signal is visible as a distortion of the topology of the tree constructed from the whole sequence compared to trees constructed from different regions. The tree constructed from the N-terminal region of 530 aa supports a bacterial origin of the sequence, the tree constructed from the C-terminal region of 180 aa supports a fungal origin of the sequence, while the tree constructed from the full sequence shows a mixed taxonomy. All trees were constructed from the first 500 BLASTP hits of respective query sequence using the BLAST web suite (full sequence; residues 1:530; residues 531:710).

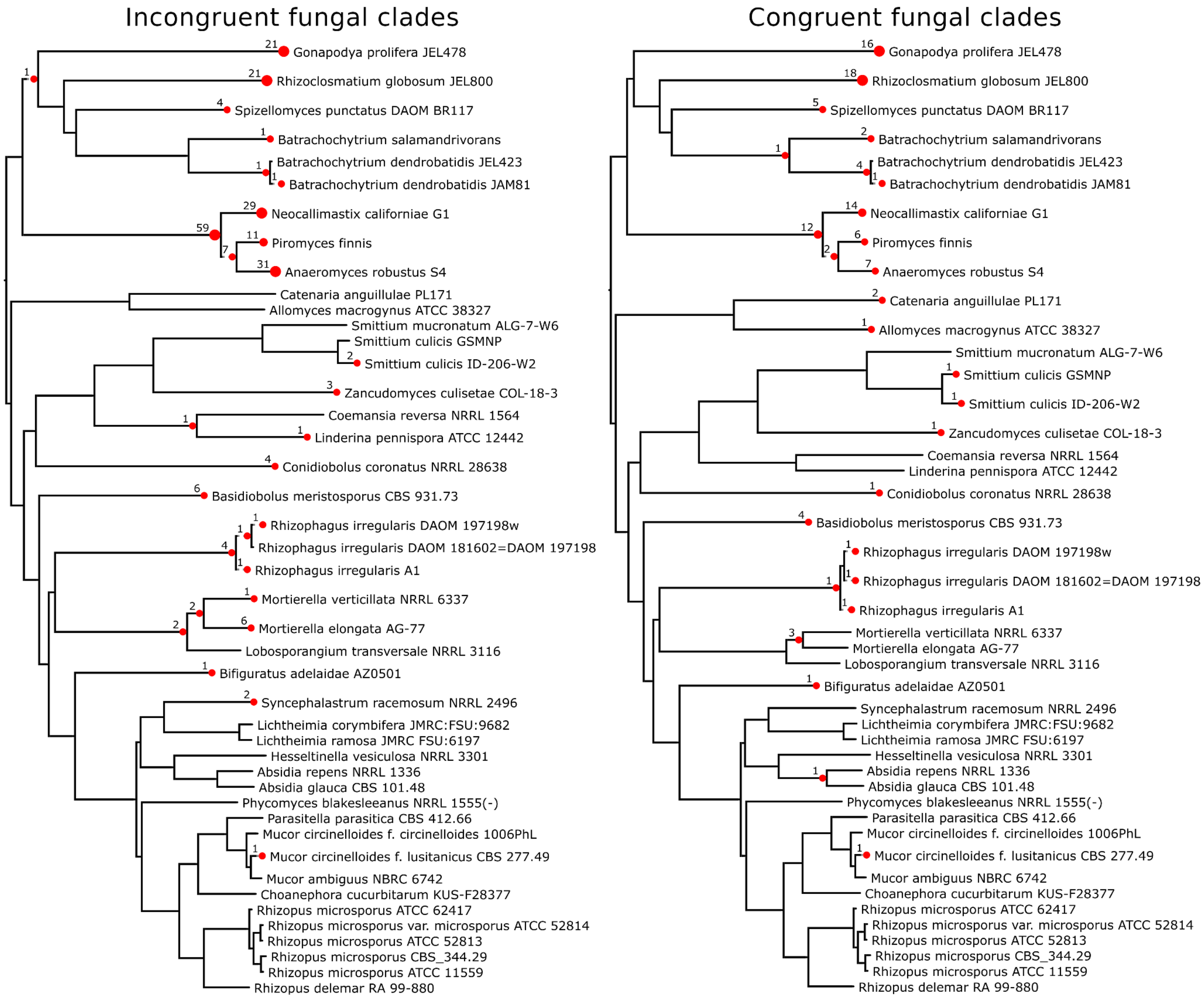


**Supplementary Fig. S7.** A comparison of the numbers of incongruent and congruent fungal clades in gene trees with predominantly non-fungal taxonomic compositions. Incongruent (displaced) clades are assumed to be a stronger indication of HGT than congruent (non-displaced) clades, as the latter may potentially also be caused by sequence homoplasy and simple vertical evolution. A high correlation of the numbers of both types of clades across the species tree of 44 EDF suggests that HGT is a more likely explanation for the congruent clades, as homoplasy would be expected to produce a uniform distribution over the tree.

### Supplementary Methods

#### Avoiding false positives in high-throughput HGT studies

There are numerous approaches to HGT detection with different scopes of applications and error rates. Phylogenetic methods, based on comparing the location of the organism of interest in the gene and the species tree of a homologous family, are considered to be the golden standard of HGT inference [1,2]. However, obtaining a gene tree is a complex procedure consisting of several steps which, in a high-throughput setting, are difficult to control and error-prone. An error in any of those steps - such as spurious homology, low-quality alignment, or imperfect tree inference - may lead to differences between the gene and the species tree and result in a false positive HGT event. During the development and testing of our pipeline, we have concluded that multiple factors can inflate the false positive rate, including:

- Relying on sequence similarity instead of phylogenetic trees. Although a useful proxy, the most similar sequence may not be the evolutionarily closest one when the molecular clock assumption is violated [3]. As a consequence, methods based solely on sequence similarity may give improper or inconsistent results [4], and methods based on phylogenies are preferred.
- Excessively long or short sequences. Both types result in poor-quality alignments and therefore in unreliable gene trees, especially when sequences of vastly different lengths are in the same cluster. Long sequences may additionally result in spurious homology, or contain regions homologous to different sets of proteins.
- Viral and environmental sequences. The placement of such sequences on the species tree is typically unknown, and their location cannot be meaningfully compared unless specific, custom-made species trees are used.
- Choice of the clustering algorithm. Since xenologs may be distantly related to their donor homolog, the clustering algorithm needs to achieve sufficient sensitivity to detect their shared ancestry, but also specificity to avoid clustering unrelated sequences. Using the high-specificity cd-hit algorithm (-c 0.7 -n 5) in our pipeline resulted in only 129 well-supported HGT events, because the clusters were overly fragmented and most of them contained proteins from less than 4 organisms, too few to conduct phylogenetic analysis. On the other hand, using the high-sensitivity mmseqs2 clustering algorithm (easy-linclust) resulted in highly unstable results, as the removal of a small subset of sequences resulted in global changes in the structure of clusters and, in consequence, different sets of HGT events. We have found that, in the context of this study, the psi-cd-hit and the orthoMCL algorithms offer the best sensitivity-to-specificity ratio.
- Low-quality alignments. Both gap-rich and highly variable sites contain little phylogenetic signal. While the former result in uncertain trees, the latter may distort the signal and the gene trees. We have found that removing highly variable sites from the alignment using the Gblocks software reduces the level of incongruence between the gene and the

species trees. On the other hand, alignment trimming algorithms that remove only the highly gapped sites and retain the highly variable ones, such as TrimAL [5], are not appropriate in the context of our study.

- Methods relying on rooted gene trees. Automatic gene tree rooting methods rely on several assumptions that are difficult to meet on the evolutionary distances encountered in large-scale studies of HGT, such as the molecular clock assumption for the midpoint method or the knowledge of duplication and transfer rates for the DTL method. When those assumptions are violated, automatic rooting methods are inaccurate and inconsistent [6,7].
- Using reconciliation algorithms with improper event weights. Any incongruence can be explained either as a gene transfer or a duplication and multiple losses, depending on the user-specified parameters which reflect the rates of those events. As we have shown in this work, transfer rates differ significantly between fungal lineages, preventing us from using reconciliation in large-scale studies. When we tested the default event weights on midpoint-rooted trees, the putative xenologs closely reflected the proteome background with respect to their PFAM domains (in particular, the regulatory WD40 and Pkinase domains were the most common), indicating that the gene trees were classified essentially at random.
- Complex gene trees with multiple fungal clades. Unless such trees are manually curated, the reconstructed evolutionary scenario is highly unstable, and the evidence for any particular transfer is less clear and unambiguous. In a large-scale study, it is difficult to determine whether multiple clades of fungi arise due to a horizontal transfer or simply an error in tree inference such as the long branch attraction.

Those considerations have led us to focus on medium-length protein sequences and gene trees with a single fungal clade displaced with respect to its location in the species tree. Our approach has some necessary limitations. Discarding long sequences has resulted in omitting some known transfers, such as a malpibaldin synthetase from *Mortierella alpina*, a 5541 amino acid-long protein (QOW41314.1). The requirement of a single fungal clade means that most detected transfers come from distantly related donors. In particular, our pipeline is unable to detect a massive transfer from a fungal parasite *Parasitella parasitica* to its host *Absidia glauca*, since both the donor and recipient are fungal species, and may have a limited sensitivity towards transfers of those animal genes which already have a fungal homolog. It is also unable to detect cases of multiple transfers of a gene, such as the seemingly independent transfers of a bacterial thymidine kinase into two microsporidian clades [8], which result in two or more fungal clades nested in the recipient’s clade. Those kinds of transfer require further studies, presumably with methods designed specifically for each scenario.

Our considerations regarding the design of a high-throughput horizontal gene transfer detection pipeline draw a parallel to the *no free lunch* theorems from mathematical optimization, which state that no single algorithm is suitable for all optimization problems [9]. In particular, studies of inter- and intra-kingdom HGT seem to have different requirements for their algorithms, just as studies of horizontal transfer of transposon or viral sequences, which have particularly complex gene trees. A single pipeline is therefore mostly suitable for a single type of HGT events, and may give spurious results when applied elsewhere.

#### Controlling long branches in gene trees

Evolutionarily distant sequences can make the topology of a phylogenetic tree unreliable despite high support values (Rodríguez-Ezpeleta et al. 2007). These include distant groups of sequences, visible in the tree as long internal branches. To avoid topological errors caused by distantly related sequences, we have adopted a strategy of iteratively splitting trees into two by cutting long branches until all remaining branches are no longer than a given threshold.

Consider an unrooted binary tree as $T = (V, E)$, i.e. an undirected acyclic graph for which each node has a degree equal either 1 or 3. With each branch *e* we have an associated branch length parameter denoted as *e*.length. Starting from a forest with a single unrooted tree $F_0 = \{T\}$, we want to find a sequence of edge cuts that results in a forest $F = \{T_1, T_2, \ldots, T_k\}$ such that each edge of each $T_i$ is shorter than a given threshold. Formally, we define cutting an edge $e = (v_1, v_2)$ of a tree *T* in forest *F* as removing the edge *e* and contracting any resulting degree-2 nodes (including summing the lengths of their adjacent branches).

Obtaining the desired forest can be accomplished by iteratively cutting a single branch with length over the threshold and updating branch lengths. Algorithm 1 shows the details of our approach. It is not difficult to show that the graph inferred by Algorithm 1 is a forest of unrooted trees. In addition, the forest does not depend on the order of long edges removals as long as $\alpha$ is fixed. From the computational complexity point of view, the algorithm can be implemented to run in $O(|T|)$ time, where $|T|$ is the size of the input tree.

---

**Algorithm 1.**
**Input:** An unrooted tree $T = (V, E)$; A branch length threshold $\alpha$.
**Output:** A forest obtained from cutting *T*'s edges, with all edges no longer than $\alpha$.
**While** there exists an edge $e \in E$ with *e*.length >= $\alpha$:
    Let $\{v, w\} = e$
    Remove *e* from *E*
    **For** *u* in $\{v, w\}$:
        **If** *u* is internal:
            Identify nodes $u_1$, $u_2$ adjacent to *u*
            Create edge $e_u = \{u_1, u_2\}$
            Set $e_u$.length = $\{u_1, u\}$.length + $\{u, u_2\}$.length
            Add $e_u$ to *E*
            Remove *u* from *V*
***Return*** (V,E)

---

#### Detecting horizontal gene transfer in unrooted gene trees

**Motivation.** The most reliable evidence for a horizontal gene transfer is based on the comparison of the topologies of two phylogenetic trees: a *gene tree*, representing the evolutionary relationship of a homologous family, and a *species tree*, representing the evolutionary relationships of species. When genetic material is exchanged between species horizontally, the resulting homologs–termed *xenologs*–start to evolve independently. This occurs after the speciation of their host species, and, in consequence, the evolutionary difference is larger for the species than for their xenologs. This leads to a characteristic *incongruence*, i.e. a difference in topology, of the gene and the species trees: the xenologs branch together in their tree, but their host species do not. An example of such incongruence is shown in Fig. S9. The analysis of tree incongruences is one of the basic tools in studying genomic events such as horizontal gene transfers and gene duplications, as well as populational effects, such as deep coalescence (an ancient separation of alleles in closely related organisms).

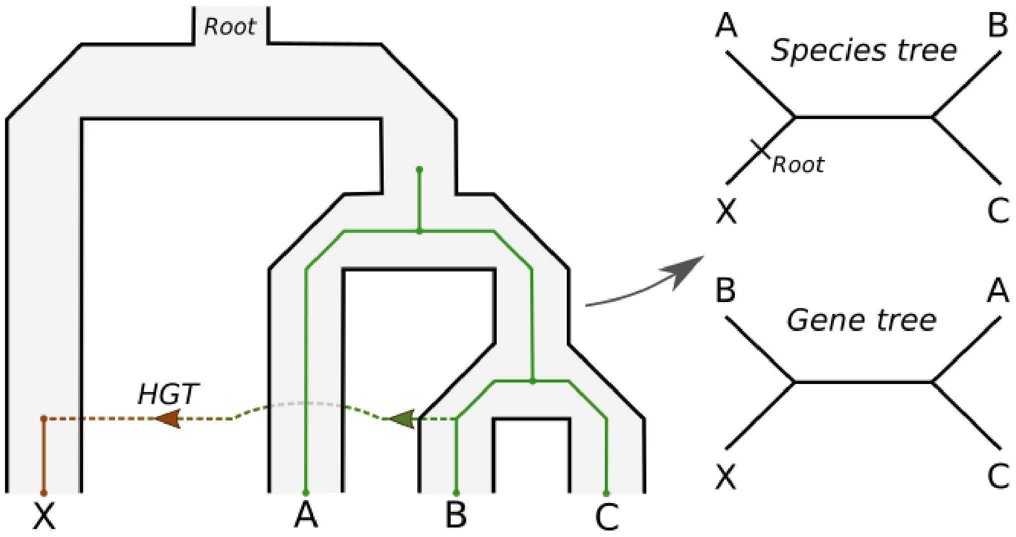


**Fig. S9:** The incongruence between trees caused by a horizontal gene transfer. The recipient clade X branches with clades {A} and {B, C} in the unrooted version of the species tree, and with clades {A, C}, {B} in the gene tree.

Arguably, the most developed methodology for studying genomic effects based on tree incongruence is the *tree reconciliation*, in which the gene tree is embedded into the species tree (with possible horizontal branches of the gene tree), in a way that minimizes a weighted sum of the numbers of different evolutionary events. With fine-tuned parameters and accurate input data, reconciliation allows to reconstruct the whole evolutionary history of a homologous family, and is an indispensable tool in detailed phylogenetic studies. On the other hand, its requirements, necessary for a detailed result, limit its applicability in a high-throughput setting. A common version of the method, the DTL model, requires the user to specify a set of weights that tell the method how likely a horizontal gene transfer is to occur in comparison to a gene duplication. However, any incongruence can be explained using only duplications or only transfers. Reliable methods of weight estimation are lacking, and it is unlikely that a single set of weights applies for all species, as the rates of transfers differ highly between them.

Another limiting requirement of tree reconciliation in the DTL model is that the most developed methods rely on the knowledge of the direction of evolution, expressed as the location of the root of the gene tree. However, the typical sequence substitution models used in maximum likelihood method of gene tree inference assume that evolution is time-reversible and result in unrooted trees. Automatic rooting methods are often inaccurate, resulting in improperly placed roots which lead to improperly reconstructed evolutionary scenarios. This adds on top of the unavoidable inaccuracies of tree inference itself. It is therefore difficult or even impossible to obtain sufficiently high-quality input data in a high-throughput setting to get a reliable sample of xenologs using tree reconciliation.

These considerations have led us to develop an alternative approach suited for detecting xenologs in a given set of genomes. In this task, we do not need to reconstruct whole scenarios, as we are only interested in testing whether a particular branch – the one adjacent to the fungal clade – represents a vertical or horizontal inheritance. We have noticed that a particular class of such transfer edges can be reliably detected in unrooted gene trees. Namely, a transfer event results in a shift of the fungal clade in the gene tree with respect to its location in the species tree. An example of such a shift is shown in Fig. S9. Therefore, the desired test can be based on a comparison of the locations of the fungal clades in the two trees.

The main difficulty in using this phenomenon to test whether a branch corresponds to a transfer is to rigorously define the notion of a location in a tree, taking into consideration natural differences between the gene and species trees. The latter often contain polytomous nodes, while the former are binary, but usually differ from the corresponding species tree in their overall topology due to other evolutionary events and tree reconstruction errors. Therefore, relying on whole topologies of the two trees to measure the location shift is not desirable.

To circumvent the topological problems associated with the different natures of both trees, we define the location shift using the label composition of the clades neighboring with the fungal clades in both trees. The general idea, formalized in this section, is that the location of a fungal clade is identical in both trees if its neighboring clades can be matched in a way that the sets of leaves of matched clades are identical, i.e. correspond to the same sets of species.

**Detecting fungal clades in gene trees.** In order to detect displaced (incongruent) fungal clades, we first need to be able to identify the clades themselves. We define an unrooted tree as a pair ($V$, $E$) of nodes and edges, where $E$ is a set of two-element sets of nodes. Let $G$ be an unrooted gene tree and $S$ be its corresponding species tree (treated as unrooted in this paragraph), so that the set of species in $S$ is the same as the set of host species of genes/proteins in $G$. We define a clade as a subtree of $G$ or $S$ that can be obtained by cutting a single edge and selecting one of the resulting subtrees. The cut edge is referred to as the clade's *attaching edge*, and the attaching edge's node belonging to the clade is referred to as the clade's *root node*.

Let *L* be the set of all fungal leaves (in either *G* or *S*). We say that a clade is induced by *L* if it contains only leaves from *L* and is maximal with respect to containment, i.e. any clade which contains it also contains leaves outside of *L*. In the gene tree, the set *L* may induce more than one clade. In order to analyze the displacement of fungal clades in *G*, we first look for all clades induced by the set of fungal leaves *L* in *G*.

Algorithm 2 shows a procedure to identify all clades induced by a given set of leaves in a given tree. It employs a boolean data structure all_in_L[*w*, *v*] which, for a directed edge (*w, v*), stores a value True if all leaves reachable from *w* through *v* are in the set of leaves *L*, and a value False if any leaf reachable from *w* through *v* is not in *L*. Algorithm 2 first identifies all directed edges whose end is a root node of some (possibly non-maximal) fungal clade. Next, it identifies *splitting edges*, i.e. edges such that there are only fungal species on one of their sides. Finally, the algorithm identifies clade attaching edges as splitting edges adjacent to some non-splitting edges. Note that this procedure works as well for non-binary trees. However, it requires each clade to have its own attaching edge. As a consequence, for non-binary species trees it may return more clades than expected if the species in *L* form a single clade in some, but not all, binarizations of the tree.

---

**Algorithm 2.**
**Input:** Unrooted tree *G* = (*V, E*); Subset of leaves *L*.
**Output:** Directed attaching edges of clades induced by *L*.

```
all_in_L = an empty map from pairs of nodes (n1, n2) to bool
clade_edges = an empty list
Function traverse_tree(w, v){
    If v is a leaf:
        If v ∈ L:
            all_in_L[w, v] = True
        Else:
            all_in_L[w, v] = False
    Else:
        Get N = a list of v's neighbors other than w
        For each node n in N:
            If all_in_L[v, n] is undefined:
                traverse_tree(v, n)
        If all_in_L[v, n] == True for all n ∈ N:
            all_in_L[w, v] = True
        Else:
            all_in_L[w, v] = False
}
For each leaf l:
    traverse_tree(l, l.parent)
For each node n:
    Get a list of n's neighbors v_1, v_2, …, v_k
    Get a boolean list b_i = all_in_L[n, v_i] for each i in 1, 2, …, k
```

**If** there exists $j$ such that $b_j$ == False:
**For** all $i$ such that $b_i$ == True:
Append ($n$, $v_i$) to clade_edges
**Return** clade_edges

---

**Detecting clade location shift.** Let $G$ be an unrooted gene tree and $S$ be its corresponding species tree, so that the set of species in $S$ is the same as the set of host species of genes/proteins in $G$. For now, we will consider a case when $G$ has a single fungal clade. Let $G_f$ denote the fungal clade in $G$ and let $S_f$ denote the fungal clade in $S$. For a given clade $C$, denote $N(C)$ the set of neighboring clades. Note that since $G$ is binary, we have $|N(G_i)| = 2$ for each $i$, but we may have $|N(S_f)| >= 2$ if (and only if) $S$ is non-binary. Let $L(T)$ be the set of species corresponding to leaves of the tree $T$.

Consider a simple case where $S$ is binary. Then, we will say that the fungal clades $G_f$ and $S_f$ have an identical location in both trees if the clades from $N(G_f)$ and $N(S_f)$ can be matched one-to-one with respect to their species composition. That is, if $N(G_f) = \{A, B\}$ and $N(S_f) = \{C, D\}$, we require that either $L(A) = L(C)$ and $L(B) = L(D)$ or $L(A) = L(D)$ and $L(B) = L(C)$. This indicates that the fungal clade has an identical branching pattern in both trees, in the sense that it branches with the same taxonomic clades regardless of their internal topology. However, $S$ may contain multiple neighboring clades, especially when highly polytomous species trees from public databases are used. In this case, we assume that a polytomy represents an unknown order of speciations, and the tree can be binarized in a way that represents the true sequence of speciation events. When comparing sets of leaf labels, a binarization of a polytomous node corresponds to merging clades from $N(S_f)$ into two sets of species. If those sets can be matched with clades from $N(G_f)$, we say that the fungal clade has an identical location in both trees.

These considerations allow us to give a simple test whether the fungal clade is displaced in $G$ with respect to its location in $S$. Let $N(G_f) = \{N^G_1, N^G_2\}$, and let $N(S_f) = \{N^S_1, \ldots, N^S_k\}$. We say that the fungal clade has the same location in $G$ and $S$ if and only if each $L(N^S_1)$ is a subset of either $L(N^G_1)$ or $L(N^G_2)$ for $i = 1, 2, \ldots, k$. This is equivalent to merging clades from $N(S_f)$ into a set corresponding to $N^G_1$ and a set corresponding to $N^G_2$. It follows that the fungal clade can be considered to be displaced if any $N^S_i$ has leaf labels present in both $N^G_i$ clades.

However, this kind of displacement can occur not only because of HGT into fungi, but also because of a HGT between species from $N(G_f)$ or a gene tree inference error. In order to differentiate between those cases, we need an additional constraint. Namely, we need to ensure that the gene tree has a branch such that, if $G_f$ was to be repositioned to that branch, it would have an identical location as $S_f$ in $S$. This means that $G$ contains a location for the fungal clade that would correspond to vertical evolution. Otherwise, $G_f$ would be considered as displaced regardless of its position in $G$. Checking for this location can be accomplished simply by ignoring fungal leaves in $G$, iterating over all edges in $G$, taking splits induced by those edges and checking if they can be matched with $N(S_f)$. Note that the lack of the location of vertical inheritance does not preclude the possibility of HGT, but our method is not suitable to detect such cases.

Consider now the case when fungi induce multiple clades in *G*, denoted $G_1, \ldots, G_m$. Then, we process each clade independently of the other, ignoring other fungal species. Namely, for each *i* we take the neighboring clades $N(G_i)$, remove any fungal species from those clades, and compare them with $N(S_f)$ (which, by definition, don't contain any fungal species). Note that checking for the location of vertical inheritance in *G* does not depend on the number of fungal clades in this tree, and can be done just once.

**Identifying the donor group**. A fungal clade with an unexpected branching pattern is indicative of a HGT into the fungi from the last common ancestor of one of the neighboring clades. In order to determine which one of the two is the donor one, we analyze their last common ancestors and the last common ancestor of the two clades combined. In the transfer scenarios analyzed in this work, the donor group branches within a larger clade of species (note that this may not be the case for ancient horizontal transfers between the ancestors of two monophyletic clades). A consequence is that one of the clades neighboring with the fungal one in the gene tree, containing species from the aforementioned larger clade, will have the same last common ancestor (in the rooted version of the species tree) as the two neighboring clades combined. For example, in Fig. S9, the last common ancestor of the {A, C} clade is identical to the last common ancestor of the two neighboring clades combined into {A, B, C}, and the last common ancestor of the {B} clade is below it. In this work, we identify the donor clade as the one for which the common ancestor is below the common ancestor of the combined clades. If both neighboring clades in the gene tree have an identical last common ancestor, we also treat it as the donor. If neither case is met, we discard the tree as either likely containing inference errors or with an evolutionary scenario too complex for an automated analysis.

6. Wade T, Rangel LT, Kundu S, Fournier GP, Bansal MS. Assessing the accuracy of

phylogenetic rooting methods on prokaryotic gene families. PLoS One. 2020;15: e0232950.

7. Lamarca AP, Schrago CG. Fast speciations and slow genes: uncovering the root of living canids. Biological Journal of the Linnean Society. 2020. pp. 492–504. doi:10.1093/biolinnean/blz181

8. Alexander WG, Wisecaver JH, Rokas A, Hittinger CT. Horizontally acquired genes in early-diverging pathogenic fungi enable the use of host nucleosides and nucleotides. Proc Natl Acad Sci U S A. 2016;113: 4116–4121.

9. David H. Wolpert and William G. Macready, “No Free Lunch Theorems for Optimization” IEEE Transactions on Evolutionary Computation Vol.1 No.1 pp.67-83,1997. Journal of Japan Society for Fuzzy Theory and Systems. 1997. pp. 696–697. doi:10.3156/jfuzzy.9.5_696
